# LAG3+ CD8+ T cell Subset Drives HR+/HER2− Breast Cancer Reduction in Bispecific Antibody Armed Activated T Cell Therapy

**DOI:** 10.1101/2025.01.04.631323

**Authors:** Robert Weldon Barnes, Archana Thakur, Suna Onengut-Gumuscu, Lawrence G. Lum, Sepideh Dolatshahi

**Affiliations:** Department of Biomedical Engineering, University of Virginia (UVA) School of Medicine, Charlottesville, Virginia 22908; University of Virginia Cancer Center, Charlottesville, Virginia, USA; Division of Hematology & Oncology, Department of Medicine, UVA School of Medicine; Department of Genome Sciences, University of Virginia School of Medicine; Beirne B. Carter Center for Immunology Research, UVA School of Medicine, Charlottesville, Virginia 22908

**Keywords:** Bispecific antibody armed T cell, Checkpoint Inhibitors, Systems Immunology, Therapeutic synergy, Mechanistic Modeling, Rational Design of Combination Therapies, LAG3, TIGIT, HR+/HER2− Breast Cancer

## Abstract

Tumor clearance by T cells is impaired by insufficient tumor antigen recognition, insufficient tumor infiltration, and the immunosuppressive tumor microenvironment (TME). Although targeted T cell therapy circumvents failures in tumor antigen recognition, suppression by the TME and failure to infiltrate the tumor can hinder tumor clearance. Checkpoint inhibitors (CPI) promise to reverse T cell suppression and can be combined with bispecific antibody armed T cell (BATs) therapy to improve clinical outcomes. We hypothesize that adoptively transferred T cell function may be improved by the addition of CPI if the inhibitory pathway is functionally active. This study develops a kinetic-dynamic model of killing of hormone receptor-positive (HR+) breast cancer cells mediated by BATs using single-cell transcriptomic and temporal protein data to identify T cell phenotypes and quantify inhibitory receptor expression. LAG3, PD-1, and TIGIT were identified as inhibitory receptors expressed by cytotoxic effector CD8 BATs upon exposure to HR+ breast cancer cell lines. These data were combined with real-time tumor cytotoxicity data in a multivariate statistical analysis framework to predict the relevant contributions of T cells expressing each receptor to tumor reduction. A mechanistic kinetic-dynamic mathematical model was developed and parametrized using protein expression and cytotoxicity data for in silico validation of the findings of the multivariate statistical analysis. The model corroborated the predictions of the multivariate statistical analysis which identified LAG3+ BATs as the primary effectors, while TIGIT expression dampened cytotoxic function. These results inform CPI selection for BATs combination therapy and provide a framework to maximize BATs anti-tumor function.

**What is already known on this topic:** Bispecific antibody armed T cell (BATs) therapies are adoptive T cell therapies that can effectively reroute T cell cytotoxicity toward cancerous cells, but lack consistent and durable anti-tumor responses. Checkpoint proteins expressed on the surface of activated T cells dampen immune responses and can be overstimulated in solid tumors to hamper tumor clearance by T cells. Checkpoint inhibitor drugs can improve T cell anti-tumor response by blocking checkpoint protein signaling but are only effective if the targeted checkpoint protein is expressed on the T cell and activated in the tumor microenvironment, highlighting an opportunity to enhance BAT efficacy by combining treatment with synergistic CPI.

**What this study adds:** This study characterizes dynamic, time-resolved patterns in checkpoint protein expression by breast cancer-targeting adoptive T cells and predicts the significance of high-prevalence checkpoint proteins on T cell function. It also demonstrates the use of multivariate statistic and mathematical modeling toward rational design of targets and timing strategies for synergistic combination therapies.

**How this study might affect research, practice, or policy:** The output of this study provides justification for therapeutic strategies combining adoptive T cell therapies with checkpoint inhibitor drugs targeting TIGIT and LAG3 as a means of improving patient responses in HER2−/HR+ breast cancers.

## INTRODUCTION

HR+/HER2− breast cancers represent 68% of all breast cancer diagnoses(1). Targeted adoptive T cell therapies offer an effective approach to elicit T cell immune responses against tumors lacking a sufficiently recognizable tumor antigen(2–6). Bispecific antibody-armed activated T-cell (BATs) therapies utilize activated T cells armed with bispecific antibodies (BiAb) targeting CD3 and a tumor antigen to redirect non-MHC restricted T cell cytotoxicity toward high and low antigen expressing cancer cells(6, 7). This strategy has shown encouraging results in reducing tumor burden, improving quality of life, and improving median overall survival (OS) in patients with stage IV breast(8, 9), prostate(10), and pancreatic(5) cancers and neuroblastoma(11).

In a phase I trial involving 17 heavily pretreated HER2−/HR+ metastatic breast cancer (MBC) patients, HER2 BATs therapy achieved a median OS of 27.4 months with 59% patients exhibiting stable disease at 14.5 weeks(12). Similarly, a phase II trial with 24 HER2−/HR+ MBC patients reported a decrease in tumor markers in 56.5% of patients and a median OS of 15.2 months following HER2 BATs infusions(13). Despite these successes, the lack of consistent and durable anti-tumor responses highlights an opportunity to enhance BAT efficacy by combining treatment with synergistic T cell targeting therapies.

Checkpoint inhibitors improve T cell function against solid tumors by preventing inhibitory signaling. While CPIs have shown efficacy in some breast cancers, response rates in HER2−/HR+ patients remain disappointingly low (5-12%) compared to HER2+ or TNBC patients(14–16). CPI efficacy depends on the presence of the target protein and its complimentary ligands or receptors, suggesting that identifying reliable CPI targets in HER2−/HR+ tumors could improve therapeutic outcomes (17, 18). By characterizing inhibitory receptor expression dynamics and BATs function, this study investigates whether the timing and selection of CPI targets can provide an opportunity for the synergistic use of CPIs with BATs therapies for HR+/HER2− patients.

Computational modeling integrates diverse measurements to provide insights into how cellular changes affect therapeutic outcomes. Network modeling has revealed therapeutically targetable gene expression mechanisms governing T cell fate selection, function, and exhaustion(20–23), but lacks temporal resolution and cannot account for repeated T cell stimulation in targeted cell therapies. Existing mechanistic models of adoptive T cell therapies have described the emergence of T cell functional states(21–23), but do not address therapeutic dysfunction.

To address this gap in knowledge, we developed a dynamic ordinary differential equation (ODE) model that incorporates BATs inhibitory receptor expression profiles. This model predicts the cytotoxic function of each BATs subset and can serve as an *in-silico* testbed to optimize the synergistic use of CPI therapies by targeting the most functional receptors at their peak expression time.

Using single-cell transcriptomics and temporal protein and gene expression data, we characterized T cell phenotype emergence and receptor expression dynamics. These data, combined with tumor cytotoxicity profiles, were analyzed using a multivariate statistical framework to predict the relative contributions of BATs subspecies to tumor cytotoxicity.

Our ODE model of BATs-mediated tumor killing was constructed from matched temporal protein expression and cytotoxicity data to validate our multivariate analysis findings. Both our multivariate statistical analysis and kinetic-dynamic model identified the LAG3+ BATs subset as a key driver of tumor reduction across multiple donors and cancer cell lines. These results suggest targeting LAG3 in BATs-CPI combinatorial therapies could significantly improve therapeutic outcomes in HR+/HER2− breast cancer patients.

## MATERIALS AND METHODS

### Target cell coculture

MCF-7, CAMA1, and T47D cells were acquired from ATCC. Cells were thawed in the media suggested by the manufacturer. MCF-7 cells were passaged in cRPMI (RPMI1640 + 10%FBS). CAMA1 and T47D cells were gradually acclimated to cRPMI by subsequent passages in mixed (RPMI/native) media with gradually increasing RPMI proportions. Cells were passaged every 2-3 days to avoid confluence.

### ATC generation

Activated T cells (ATC) were generated using T cells collected from donor PBMC. Blood was collected from normal donors and PBMC were obtained by Ficoll-Paque density gradient separation, counted, suspended in RPMI supplemented with 100 U/mL IL-2 and 20 ng/mL OKT3 to activate T cells and stimulate proliferation(24). T cells were expanded by refreshing the cultures with RPMI containing 100 U of IL-2/mL every 2-3 days for 14 days. The expanded T cells were harvested and cryopreserved at 100e^6^ viable cells (vc)/mL at −160°C in LN2 until use. ATC used in this study were collected from three donors. Donor information was anonymized prior to the experiment with the exception of sex. Two donors were female and one donor was male.

### HER2-targeting Bispecific Antibody Preparation

The HER2-targeting bispecific antibody (HER2bi) was generated by chemical heteroconjugation of OKT3, a murine IgG2a anti-CD3 monoclonal antibody (Miltenyi Biotec), and Herceptin, a humanized anti-HER2 IgG1 monoclonal antibody, using sulfo-succimidyl 4-(N-Maleimidomethyl) cyclohexane-1-carboxylate (Sulfo-SMCC) and Traut’s reagents as described previously(25). An SDS-PAGE analysis of the HER2bi used in this study is shown in **Fig S1**.

### BATs preparation

BATs were prepared as described(26). Briefly, ATC were thawed at 37°C, resuspended in RPMI, and washed 2x in RPMI. Washed ATC were resuspended in RPMI + 50 ng/10^6^ vc anti-HER2x anti-CD3 bispecific antibody (HER2Bi) and rocked on a rocking platform for 15 minutes to allow for HER2Bi to attach to ATC to generate HER2 BATs(27). HER2 BATs were washed 3x to remove excess biab and resuspended at 10^6^ vc/mL. A pictorial description of BAT generation can be found in **Fig S2**.

### Single-cell cellular indexing of transcriptomes and epitopes by sequencing (CITE-seq) Analysis

ATC from 3 donors were thawed and split into two groups per donor. One group from each donor was armed as described above to generate BATs. Each group was cocultured for 1 hours with MCF7 tumor cells at a 10:1 BAT:MCF7 ratio. BATs were then collected and measured using 10X Genomics CITE-seq technology(27). TotalSeq^TM^-B oligonucleotide-labeled antibodies for CD45RA, CD3, CD4, CD8, CD69, and CD25 were included in the CITE-seq panel, and cells were stained following the manufacturer’s protocol (BioLegend). CITE-seq libraries were processed in parallel following 10X Genomics Single Cell 3’ protocol and sequenced on an Illumina NextSeq 500.

Raw protein tag and transcriptomic data from each donor were integrated into a Seurat object using the Seurat 5.0 R package(28). Each object was filtered to remove cells with < 500 features or > 2500 features, as well as those with < 1% or > 5% mitochondrial DNA. Data within each object was normalized and the 3000 most variable features were identified. All 6 objects were integrated into a single object using Seurat FindIntegrationAnchors and IntegrateData functions. Data was then scaled, and a principal component analysis (PCA) was conducted to identify the top 100 principal components (PCs). A K nearest-neighbor graph was constructed using the top 25 PCs and used to identify cell clusters. A uniform manifold approximation and projection (UMAP) plot for visualization was created using these clusters. Differential gene expression was used to identify T cell clusters. Briefly, a Wilcoxon Ranked Sum test was run using Seurat’s FindAllMarkers function. Genes strongly differentiating each cluster were cross-referenced with T cell literature to identify the phenotype in question.

### Real-time cell analysis (RTCA) measurements

The xCelligence RTCA monitor was used to measure tumor cell death across the course of the assay. xCelligence plates were zeroed with RPMI prior to seeding(29). In brief, 2e4 vc target breast cancer cell lines cultured in RPMI1640 supplemented with 10% FBS were added to each xCelligence well and allowed to adhere. Incubator conditions were maintained at 37 °C and 5% CO2. After 24 hours, BATs were added to each well at predefined ratios between 0.125 – 2 BATs:Tumor. As a surrogate for cytotoxicity, conductivity measurements were recorded continuously for 120h.

### BATs coculture preparation

Time-course samples were prepared by coculturing BATs and tumor cells at an effector: target ratio of 1:1. Flat bottom 24 well plates were seeded with MCF-7 cells in RPMI at 1.2e5 vc/mL and incubated for 24 hours. After 24h, BATs were added at a E:T as determined by target cell counts from a sample plate. Bat-tumor cocultures were incubated for up to 72 hours. Samples were collected from wells at 4, 24, 48, and 72 hours. Collected samples were used for flow cytometry readings.

### Checkpoint inhibitor testing

To test the effects of checkpoint inhibitors on BAT function, RTCA plates were seeded with 2x 10^4^ vc/well MCF7 cells, allowed to adhere, and grown for 24 hours. Conditions and media were identical to those described above. BATs were then mixed with either 10.0, 3.2, 1.0, 0.32, 0.1, 0.032, or 0 ug/mL Tiragolumab biosimilar InVivoSIM anti-human TIGIT or Relatlimab biosimilar InVivoSIM anti-human LAG3, incubated for 30 minutes at 37C, then transferred at a 1:1 BAT:Tumor ratio into corresponding RTCA wells. BAT-tumor cocultures were incubated for up to 72 hours. An identical setup in 96 well plates was performed separately to generate flow cytometry samples. Samples were collected from wells at 4, 24, 48, and 72 hours. Collected samples were used for flow cytometry readings.

### Flow Cytometry

Flow cytometry stain panels (**Table 1**) were created by mixing fluorescence-conjugated antibody stains targeting the ligands of interest. The following panels were generated: Inhibitory (CD45, CD4, CD8, TIGIT, PD-1, LAG3, TIM-3, CTLA, BTLA), Stimulatory (CD45, CD4, CD8, 4-1BB, CD27, OX40, GITR, HVEM, LIGHT, ICOS), and Coexpression (CD45, CD4, CD8, 4-1BB, 4-1BBL, OX40, TIGIT, PD-1, LAG3). Cells were resuspended, counted for viability, stained in 4 °C DPBS, and fixed before fluorescent intensity readings were acquired on the NovoCyte 3005 3 laser cytometer. Collected data was compensated and analyzed using FlowJo flow cytometry data analysis software. Cells were identified using size and complexity characteristics identified from a T cell-only control sample. Live cells were identified using LIVE/DEAD Fixable Violet Dead Cell stain kit. T cells were further separated from tumor cells by either CD3 or CD45 expression. T cells were further subset using anti-CD4 and anti-CD8 antibodies. A sample flow gating strategy is shown in **Fig S3**. The MFIs of each target stain can be seen in **Fig S4**. Prevalence of each receptor was calculated by percentage expression within CD4 or CD8 subsets.

**Table 1:**
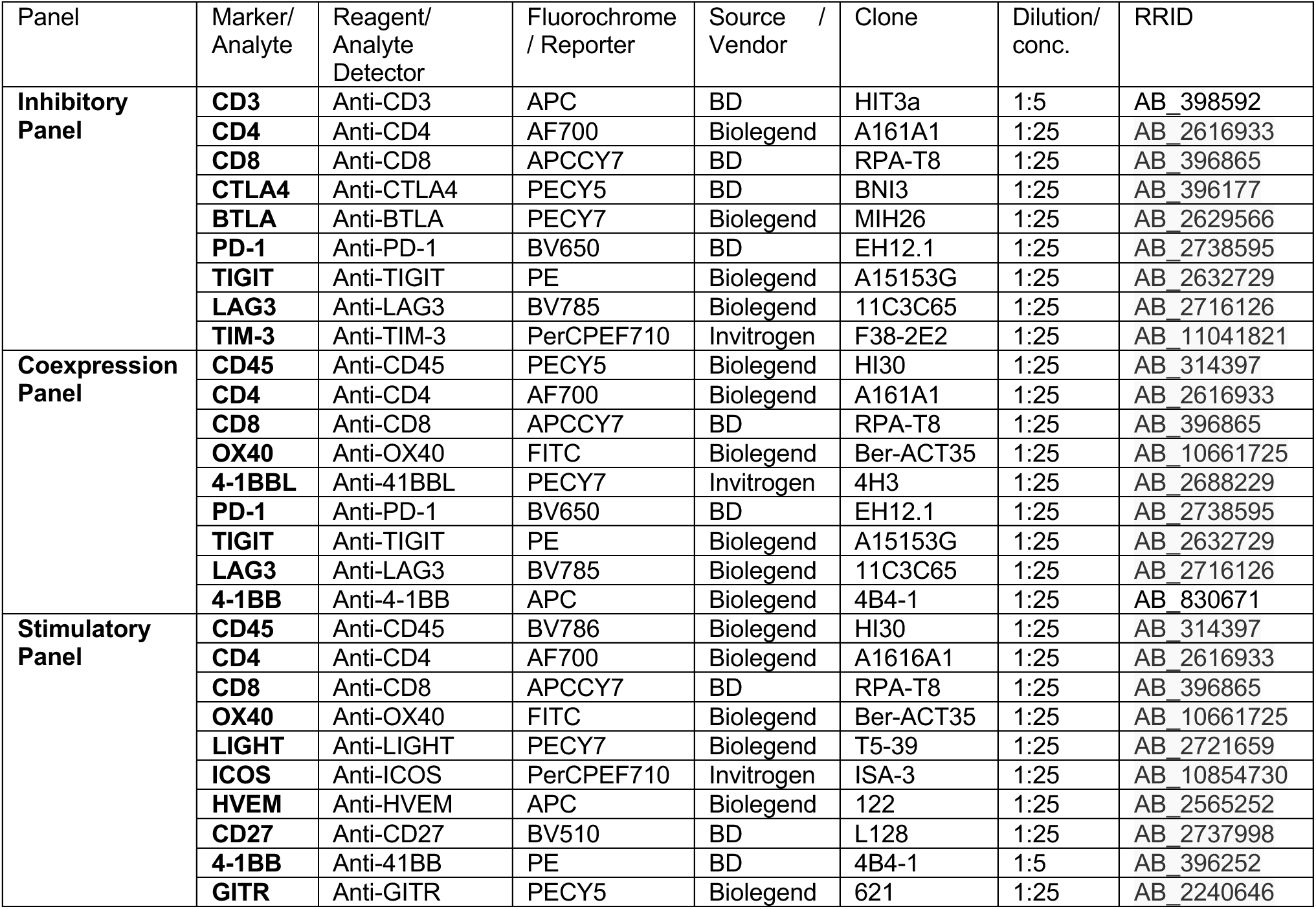
Targets for fluorescent antibodies included in each flow cytometry panel.

### OPLS-R analysis

An orthogonalized partial least squares (OPLS)-regression model was built to predict the impact of BATs receptor expression on tumor reduction. BATs were binned into distinct groups by their expression of CD8 and CD4, and again by their expression of PD-1, TIGIT, and/or LAG3. Triplicate counts were averaged within donor and against target cell line. These counts were inputted to predict the changes in tumor cell count and the rate of change in tumor counts as measured by RTCA. Separate models were created for the 3-donor study and the 3-target study.

Inputs for tumor counts or rate of change compared against BATs counts were taken at a set time interval from the time of BATs collection. Models for each study were created with 0, 1, 4, 12, and 24-hour intervals from the time of BATs collection to increase the robustness of model predictions. The variable importance in projection (VIP) scores for each BATs subset were calculated(30), ranked, and artificially oriented in the direction of their loadings on latent variable 1 (LV1). The directionality of groups with VIP scores >1 was used to predict the positive or negative impact on tumor growth for each receptor/ligand.

### ODE model construction and testing

An ODE model was constructed using mass action equations to describe changes in tumor cell count. The following equation was used to describe tumor cell count.

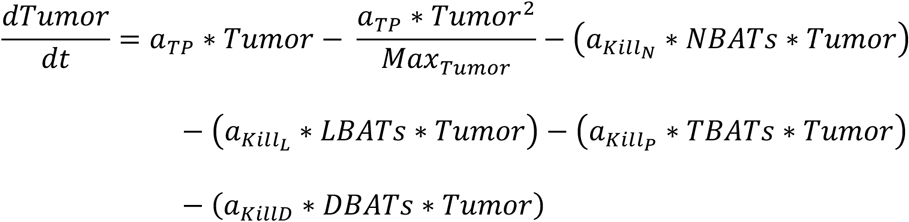

The first two terms represent tumor growth in the absence of BATs, and the next four negative terms represent the contribution of each BAT subset to the rate of tumor cell clearance. Tumor growth rate was assumed to be sigmoidal, and was fit to data collected from the control 0:1 BATs:Tumor condition during the RTCA experiment. NBATs (LAG3− TIGIT− CD8s), TBATs (LAG3− TIGIT+ CD8s), LBATs (LAG3+ TIGIT− CD8s), and DBATs (LAG3+ TIGIT+ CD8s) rate of change were determined by taking the derivative of spline functions fit to the flow cytometry cell counts collected at 0, 4, 24, 48, and 72 hours.

The CaliPro parameter optimization package was used to determine the potential ranges and optimal values for each species rate parameter (31). CaliPro optimizes unknown model parameters by creating a parameter space using Latin Hypercube Sampling within parameter-specific ranges set by the user. Each parameter was tested by software, and parameter sets that meet a user-defined set of criteria are used to generate a new LHS parameter space for a second iteration. This procedure continued until a large enough portion of parameters met the acceptance criteria. The final parameter space contains a refined range of parameters which best describe the system in question. Acceptable parameter spaces were those that generated tumor counts within +/− 10% of the measured values at 0, 4, 12, 24, 36, 48, 60, and 72 hours.

## RESULTS

### Bispecific antibody-driven interactions generate highly cytotoxic LAG3+ effector T cell populations after 1 hour exposure to target cells

HER2 BATs were generated by arming activated T cells from donor PBMC with a HER2-targeting bispecific antibody (**Fig S1**) (Materials and Methods) and cocultured with HR+ breast cancer cells. To identify phenotypic changes in T cells driven by the engagement of T cell receptor (TCR) and tumor antigen mediated by the BiAb, we profiled transcriptional and protein expression changes in HER2 BATs and unarmed ATC after exposure to tumor cells using CITE-seq (**Fig 1A**). The matched protein and RNA data were integrated and pooled, and an unsupervised graph-based clustering method was applied to generate 12 cell clusters (**Fig 1B**). CD4 and CD8 protein tags were used to identify major T cell populations, while CD69 and CD45RA protein tags helped differentiate between resting, dividing, and activated T cell subsets (**Fig 1B, Fig S5**). These 12 cell subsets exhibited varying transcriptomic expression profiles of cytotoxic factors, inhibitory receptors, costimulatory receptors, and relevant transcription factors (**Fig 1D**). Wilcoxon ranked-sum testing was used to identify marker genes distinguishing each group (**Fig 1C**) to assist in applying known T cell phenotypes.

**Figure 1:**
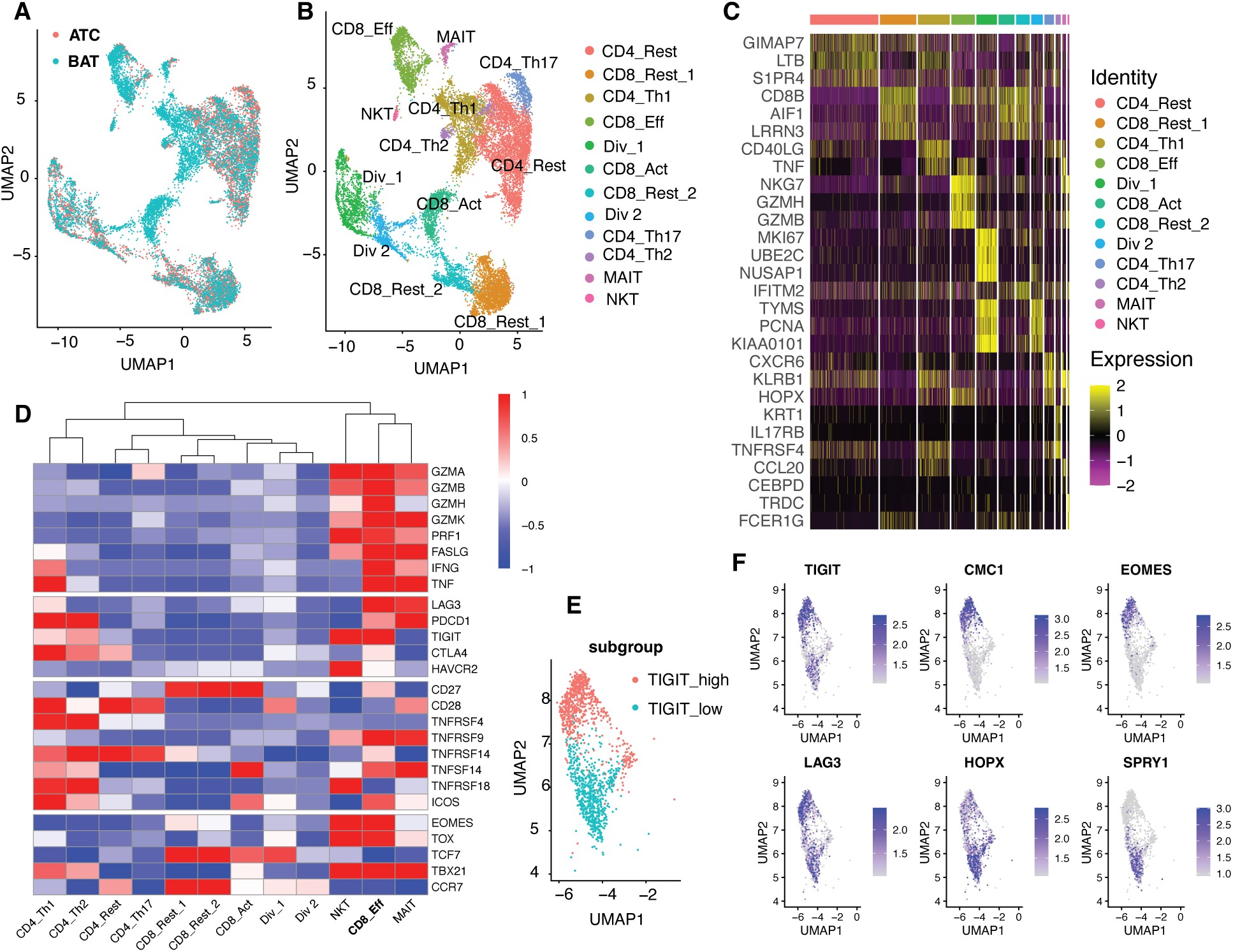
A: UMAP plot of CITE-seq data labeled by bispecific antibody arming. ATC: unarmed Activated T Cells, BAT: Bispecific antibody-armed Activated T cells B: A UMAP plot of T cells clustered by similarity in gene expression. Labels were chosen based on expression of significantly upregulated marker genes. CD4_Rest = Resting CD4 T cells; CD8_Rest_1 = Group 1 Resting CD8 T cells; CD4_Th1 = Th1 CD4 T cells; CD8_Eff = Effector CD8 T cells; Div_1 = Group 1 Dividing T cells; CD8_Act = Activated CD8 T cells; CD8_Rest_2 = Group 2 Resting CD8 T cells; Div_2 = Group 2 Dividing T cells; CD4_Th17 = Th17 CD4 T cells; CD4_Th2 = Th2 CD4 T cells; MAIT = Mucosal Associated Invariant T cells; NKT = NK-like T cells C: Heatmap of most differentially expressed genes as determined by a Wilcoxon ranked sum test. Genes shown most strongly differentiate each group from all other groups and were used to determine the labels in 1B. D: A heat map of average gene expression scaled across groups. Genes displayed are involved in cytotoxicity (top), T cell inhibition (second top), T cell costimulation (second bottom), and T cell maturation and differentiation (bottom). E: A re-clustered UMAP of effector CD8 T cells. F: Differentially expressed genes between the two CD8_Eff clusters.

The presence of the BiAb drove phenotypic changes in both CD4 and CD8 T cells (**Fig 1A, B**). Activated CD4 BATs, indicated by CD69 protein expression, switched primarily to a pro-inflammatory T helper 1 (Th1) phenotype (**Fig 1C**, **Fig S5**). A smaller subset of activated CD4 BATs was observed to adopt a humoral Th2 phenotype (**Fig 1B**), which not only drives B cells to produce immunoglobulins but may also facilitate tumor progression(32). The Th1 population was comprised primarily of BATs, which accounted for 82% of the population, while the Th2 population was evenly split between BATs (46%) and ATC (54%). CD8 BATs displayed progressive activation: a mid-activation group, labeled CD8_Act, was CD45RA-CD69+, while a fully activated effector group, labeled CD8_Eff, was CD45RA+CD69+. Additionally, two smaller BATs groups, an NK-like (NKT) group marked by KLRD1(33) and FCER1G(34) and mucosal-associated invariant (MAIT) group marked by KLRB1(35) and NCR3(36), were also present. The CD8_Eff group was identified as the primary cytotoxic group, as demonstrated by high expression of granzyme (GZMA, GZMB, GZMK, GZMH) and perforin (PRF) genes, as well as inflammatory cytokine (IFN-γ, TNF-α) and FAS ligand expression (**Fig 1D**). The CD8_Eff group was also enriched in EOMES and TBX21, two transcriptional regulators associated with T cell maturation, terminal differentiation, and effector function(37), further indicating the role of this group as an effector T cell (**Fig 1D**). TCF7, a transcriptional regulator indicating a pre-antigen encounter T cell state(38), was downregulated in the CD8_Eff group, while TOX, a gene associated with T cell effector function and exhaustion(39), was upregulated (**Fig 1D**).

Strong cytotoxic potential was observed to correlate with targetable inhibitory and costimulatory receptor expression (**Fig 1D**). The cosignaling receptors LAG3 and TIGIT, which directly modulate BATs cytotoxicity, were enriched in the CD8_Eff group (**Fig 1E**). While expression of PD-1 (PDCD1), TIM3 (HAVCR2), and CTLA4 was detected in the CD8_Eff group (**Fig 1D**), their levels were low and not significantly different from those in the total cell population.

Costimulatory receptors 4-1BB (TNFRSF9), ICOS, CD27, HVEM (TNFRSF14), and LIGHT (TNFSF14) were observed in the CD8_Eff group (**Fig 1D**). LIGHT and ICOS expression were upregulated in all activated BATs groups but were not significantly upregulated in the CD8_Eff group (**Fig 1D**). CD27 was present but downregulated compared to other CD8 groups (**Fig 1D**), indicating terminal effector differentiation(40). Only 4-1BB expression was significantly upregulated in the CD8_Eff group (See

## Discussion

While LAG3 expression was observed throughout the CD8_Eff group, TIGIT expression was concentrated within a subset of these cells (**Fig 1E, F**). Comparison between TIGIT_High_ and TIGIT_Low_ CD8_Eff cells reveals distinct expression of regulatory genes associated with T cell effector differentiation. TIGIT_High_ CD8_Effs are enriched for EOMES and CMC1, while TIGIT_Low_ CD8_Effs are enriched for HOPX and SPRY1. All four genes are involved in T cell effector differentiation(37, 41–43), but the absence of overlap may indicate a potential divergence in function between TIGIT_High_ and TIGIT_Low_ CD8_Eff cells. These expression patterns may indicate that highly cytotoxic CD8 BATs are more strongly regulated by TIGIT or LAG3 after initial interaction with breast cancer tumor cells and might benefit from CPI therapies targeting these receptors. Likewise, 4-1BB agonism may provide costimulatory signals preferentially to a highly cytotoxic subpopulation.

### Temporal profiling reveals distinct dynamics of costimulatory and inhibitory receptor expression following tumor exposure

The emergence of therapeutically targetable cosignaling receptors as marker genes for highly cytotoxic effector cells after 1 hour of exposure suggests promising potential for effective combinatorial BATs therapies. To determine whether the observed expression patterns of TIGIT, LAG3 and 4-1BB from CITE-seq data persist beyond 1 hour of stimulation and whether transcriptomic expression patterns translate to expression of surface proteins we analyzed surface receptor expression on BATs cocultured with the breast cancer cell lines. The coexpression dynamics of inhibitory and stimulatory receptors on BATs were profiled over a 72-hour coculture with HR+ tumor cells (**Fig 2A**). Flow cytometry panels used to profile receptor expression were designed based on gene expression values observed in **Fig 1D** and are detailed in **Table 1**.

**Figure 2:**
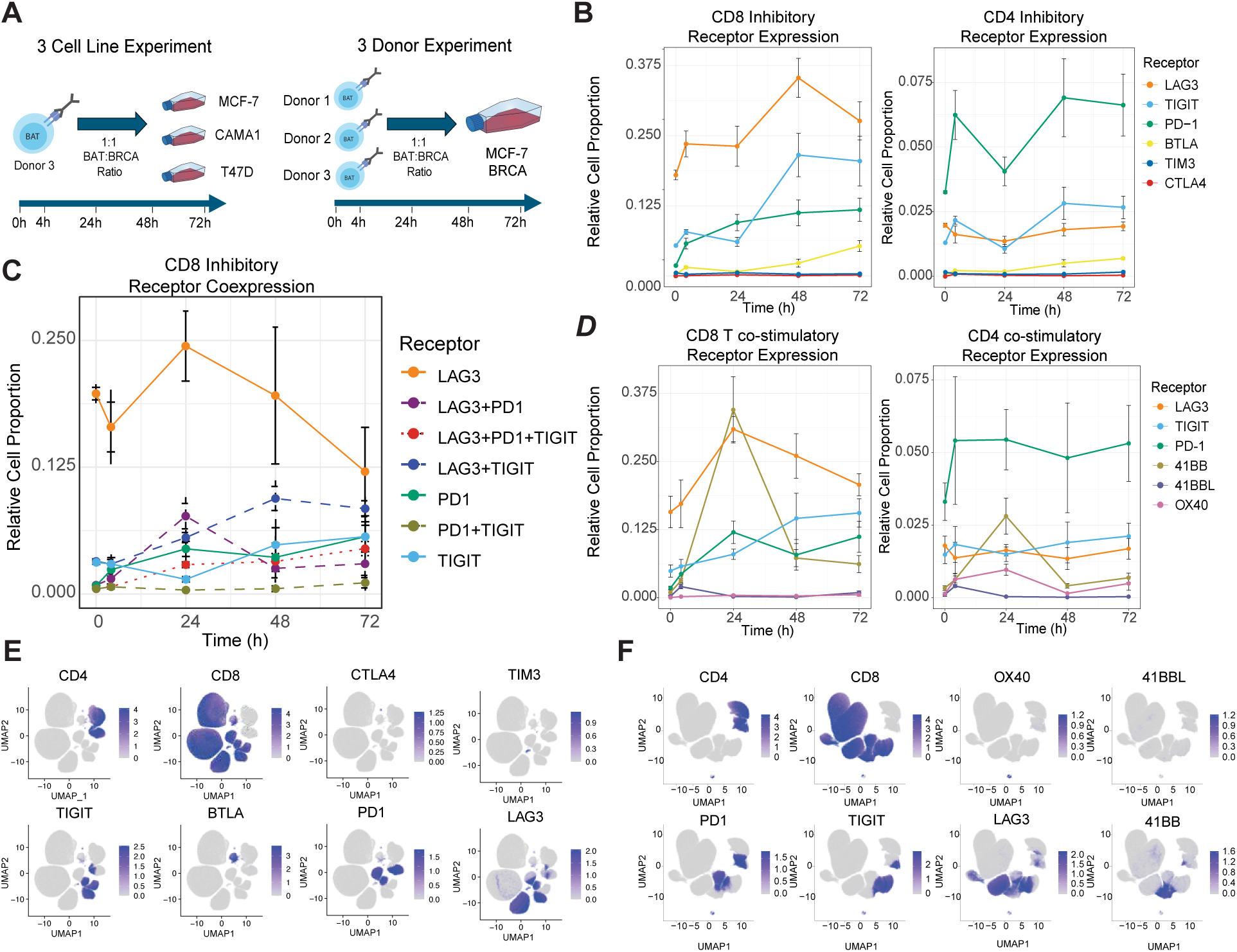
A: Diagram illustrating the setup and data collection for the two sets of flow cytometry experiments performed. Left Panel - BATs generated from Donor 1 challenged against MCF-7, CAMA1, and T47D breast cancer cell lines. Samples collected at 0, 4, 24, 48, and 72 hours and designated ‘3 Cell Line Experiment’. Right Panel - BATs generated from Donors 1, 2, and 3 challenged separately against MCF-7 breast cancer cell line. Samples collected at 0, 4, 24, 48, and 72 hours and designated ‘3 Donor Experiment’. B: Inhibitory panel receptor expression on CD8 (left panel) and CD4 (right panel) BATs. Data shown was collected from the 3 Cell Line experiment and is presented as the mean value across all cell lines. Error bars indicate standard error within each data group. Each line represents the relative proportion of the cells that express a marker regardless of other marker expression. C: Relative cell proportion of CD8 cells expressing LAG3, TIGIT, PD-1, or a combination thereof. Cells were binned specificly by their expression or combination of expression of LAG3, TIGIT, and PD-1. Unlike Panels B and D, there is no overlap between groups containing a common ligand. For example, the TIGIT+ CD8 T cell subset does not express PD-1 or LAG3. Samples collected at 0, 4, 24,48, and 72 hours. Data represents mean value across 3 Cell Line experiment as collected in the coexpression panel. Error bars represent the range within each sample timepoint. D: Coexpression panel receptor expression on CD8 (left panel) and CD4 (right panel) BATs. Data shown was collected from the 3 Cell Line experiment and is presented as the mean value across all cell lines. Error bars indicate standard error within each data group. Each line represents the relative proportion of the cells that express a marker and can have other markers simultaneously expressed. E: UMAP of BAT cells from the 3 Cell Line experiment clustered by receptor expression as measured by flow cytometry in the inhibitory panel. F: UMAP of BAT cells from the Cell Line experiment clustered by receptor expression as measured by flow cytometry in the coexpression panel.

#### CD4 BATs preferentially express PD-1, which coexpresses with LAG3 and TIGIT

CD4 BATs expressed PD-1, TIGIT, and LAG3 during the initial 72-hour challenge (**Fig 2B**). CD4 BATs preferentially expressed PD-1, while LAG3 and TIGIT were primarily coexpressed on PD-1 expressing cells (**Fig 2E**). ICOS expression within the CD4 pool gradually increased with time, while CD27 expression dipped over 24h before rebounding (**Fig S6A**). Th1 CD4 cells expressed PD-1 significantly higher than other BATs groups in our CITE-seq data, potentially suggesting the immediate emergence of a Th1 CD4 subset (**Fig 1D**). Th1 CD4 cells expressed PD-1 significantly higher than other BATs groups in our CD4 data, potentially suggesting the immediate emergence of a Th1 CD4 subset.

#### CD8 BATs rapidly express LAG3 and 4-1BB while gradually increasing PD-1 and TIGIT expression

The inhibitory panel showed that CD8 BATs preferentially expressed LAG3 (**Fig 2 B, C**). TIGIT and PD-1 were observed to a lesser extent on CD8 T cells, and primarily coexpressed with LAG3 (**Fig 2 C, E, Fig S7**). TIM3, CTLA4, and BTLA were minimally expressed by BATs over the 72 hours duration of the experiment. In all samples, LAG3+ CD8 BATs hit a proportional and numerical high between 24 and 48 hours after coculture, before declining across the remaining assay duration. PD-1 and TIGIT expression gradually increased as the assay progressed. Interestingly, samples that maintained a higher proportion of LAG3+ CD8 T cells throughout the assay cleared a greater proportion of tumor cells cumulatively across 72 hours. A stimulatory receptor profile demonstrated CD8 BATs primarily expressed 4-1BB and ICOS, with 4-1BB spiking rapidly during the first 24h before exiting the system and ICOS gradually increasing with time. CD27, which was strongly expressed at t = 0 hours, rapidly decreased in expression during the first 24 hours, before recovering to initial expression levels during the last 48 hours of the experiment (**Fig S6A**). Costimulatory receptor expression overlap was minimal, with the exception of CD27 coexpression (**Fig S6B**). The presence of LAG3, TIGIT, and 4-1BB indicates the CD8 BATs are primarily the CD8_Eff phenotype observed in our CITE-seq data.

A coexpression panel was designed to explore overlap between costimulatory and inhibitory receptors (**Fig 2D, F**). Expression of coinhibitory receptors reflected what was observed on the inhibitory panel – LAG3 and PD1 expression dominated CD8 and CD4 BATs, respectively, with both PD-1 and TIGIT coexpressing with LAG3 at increasing frequency on CD8s (**Fig 2D, F**). PD-1 and TIGIT rarely coexpressed on CD8s in the absence of LAG3. Interestingly, 4-1BB strongly coexpressed with LAG3 on CD8s during the first 24h (**Fig 2D, F**). LAG3 and 4-1BB presence simultaneously peaked at 24 hours, and LAG3 and TIGIT remained the primary inhibitory receptors after 24 hours, (**Fig 2D, F**), which suggests that 4-1BB expression or stimulation precedes T cell activation pathways that preferentially express LAG3 and/or TIGIT.

### Multivariate statistical analysis predicts opposing effects of LAG3 and TIGIT expression on BATs-mediated tumor cell cytotoxicity rates

We hypothesized that LAG3, TIGIT, and PD-1 represent distinct anti-tumor functional phenotypes, with their expression kinetics closely correlating with changes in BATs function and phenotype. PD-1 was enriched in activated Th1 and Th2 CD4 subgroups but did not associate strongly with CD8_Eff in our CITE-seq data (**Fig 1D**). LAG3 and TIGIT correlated strongly with components of cytotoxic pathways in our CITE-seq data (**Fig 1D**) and suggests a highly cytotoxic activated effector state in BATs. However, TIGIT was only partially enriched within our CD8_Eff group and correlated with regulatory components associated with effector cell maturation (**Fig 1D**). A Partial Least Squares Regression (PLS-R) model was constructed to determine if tumor cell reduction cytotoxicity could be predicted by receptor presence (**Fig 3A, B**). Cells were binned by CD4/CD8, TIGIT, LAG3, and PD-1 expression, with cells expressing combinations of these receptors put in unique bins. The prevalence of each phenotype at each timepoint was used to predict the rate of tumor cytotoxicity measured by RTCA at 4 hours post-receptor measurement. Our model predicted 98% of the variance seen in the rate of tumor growth and reduction, and outperformed 1000 random permutations of the data (p <0.001) (**Fig 3A**). Examining the top predictors of the cytotoxicity, which were defined as those with VIP >1, we observed that CD8 BATs expressing LAG3 with or without PD-1 correlated with increased rates of cytotoxicity, while TIGIT expression on either CD8 or CD4 cells was negatively correlated with the rate of cytotoxicity (**Fig 3 B, C**). Cells expressing a combination of LAG3 and TIGIT, regardless of PD-1 expression, were not predictive. Cells expressing none of the three receptors were negatively correlated with cytotoxicity, similar to TIGIT expressing cells, potentially suggesting a pre-activated state. In summary, our data show that the expression of LAG3 on BATs is significantly associated with anti-tumor cytotoxicity, while TIGIT expression dampens BATs-driven cytotoxicity (**Fig 3C, Fig S8**). Considering these results, we predict CD8 BATs will fall into one of 4 generalized functional groups: NBATs (LAG3−/TIGIT−), LBATs (LAG3+/TIGIT−), TBATs (LAG3−/TIGIT+), and DBATs (LAG3+/TIGIT+), with exposure to tumor cells driving the transfer of BATs from a LAG3 and/or TIGIT negative group to a LAG3 and/or TIGIT positive group (**Fig 3D**).

**Figure 3:**
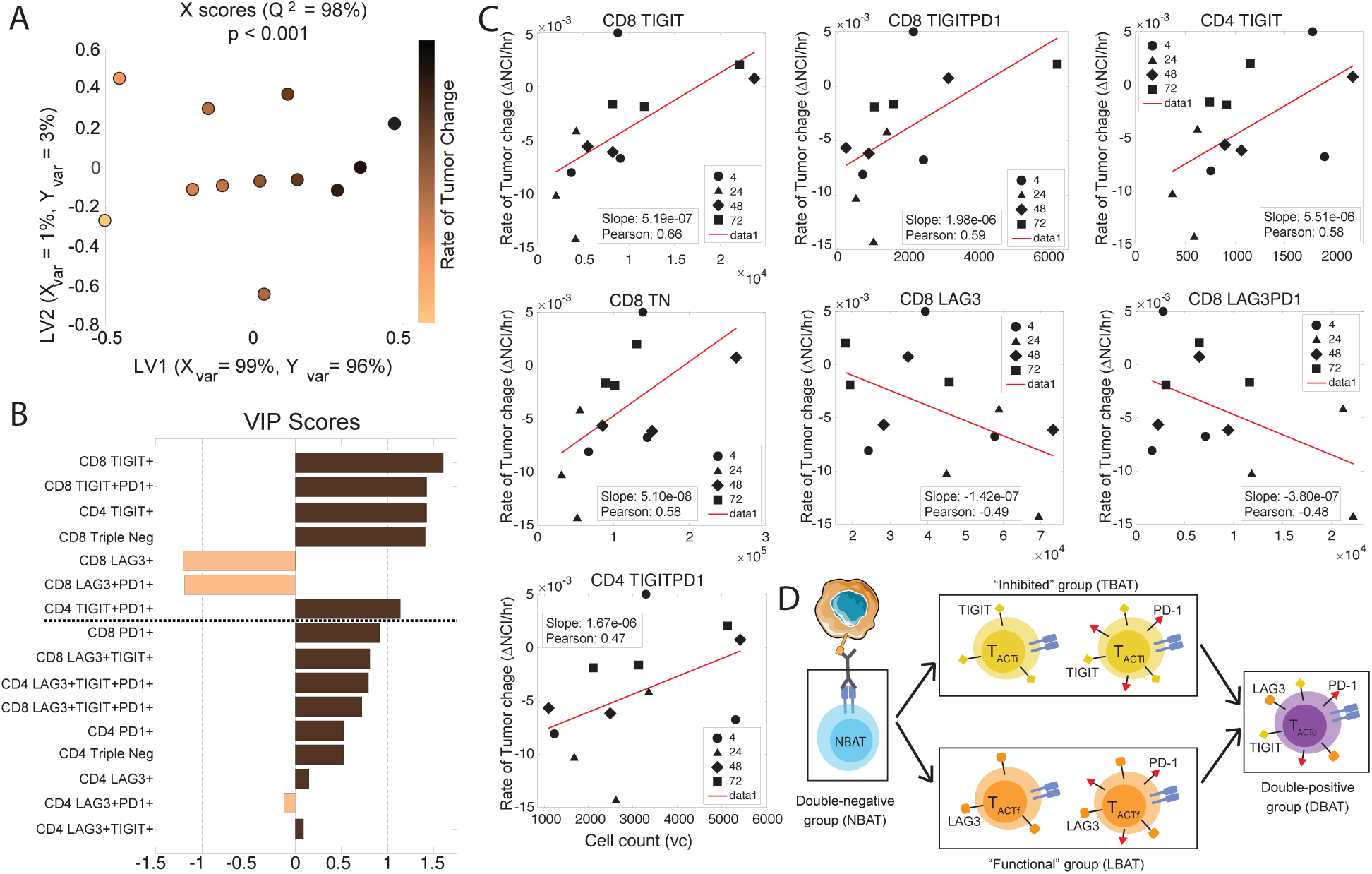
A: OPLS-R model of rate of tumor change predicted by BAT subpopulations from the 3 Cell Line experiment. BATs were binned by CD4 or CD8 expression, and further binned by expression of LAG3, TIGIT, and/or PD-1. Cells expressing more than one receptor were binned in separate categories, with cells expressing all three receptors being designated tri-positive (TP). BATs lacking receptors were designated tri-negative (TN). B: VIP scores generated by PLS-R model in Figure 3A. Subpopulation impact is designated by the magnitude of the VIP score, with scores of magnitude >1 designated as greater-than-average contribution of that cell subpopulation to tumor reduction. Subpopulations with scores colored tan are associated with reduced rate of tumor change, while subpopulations with scores colored brown are negatively associated with reduced rate of tumor change. C: Scatter plots comparing the viable cell count of BAT subpopulations (x-axis) to rate of tumor change (y-axis) for subpopulations with VIP scores >1. NCI = normalized cell index. D: Diagram of BAT developmental progression as assumed by the model. BATs are grouped as NBATs (LAG3−/TIGIT−), LBATs (LAG3+/TIGIT−), TBATs (LAG3−/TIGIT+), or DBATs (LAG3+/TIGIT+). PD-1 expression does not factor into grouping and can be present within any group. These four groups are used to govern tumor reduction in the models presented in Figures 4 and 5.

### Mechanistic modeling identifies LAG3 BATs as key drivers of tumor reduction

The PLS-R model indicates that the balance of BATs subsets defined by receptor expression plays a crucial role in tumor reduction. To more directly determine the functional impact of each subset and validate our PLS-R findings, an ODE-based dynamic model of tumor-BATs interactions was constructed (See Methods). Surviving tumor cell counts were extracted from RTCA data, while BATs subset counts were extracted from receptor expression measured by the coexpression flow cytometry panels. Given the homogeneity in CD4 population dynamics across experiments, cytotoxic CD8 cells were selected for the development of the model. Cytotoxic potential as predicted for each subset by the PLS-R model was used to define four functional groups in the kinetic-dynamic BATs-tumor model. Specifically, the PLS-R model predicted LAG3 and TIGIT expression have significant but opposing effects on the rate of tumor expansion or survival. LAG3 expression predicts higher rates of tumor reduction, while TIGIT predicts weaker tumor reduction. The expression of PD-1 was not predictive. Based on the above findings, a mechanistic model of tumor-BATs interactions was constructed consisting of tumor cells and 4 receptor-defined BATs species: LAG3−/TIGIT− BATs (NBATs), LAG3+/TIGIT− BATs (LBATs), LAG3−/TIGIT+ BATs (TBATs), and LAG3+/TIGIT+ BATs (DBATs) (**Fig 4A**).

**Figure 4:**
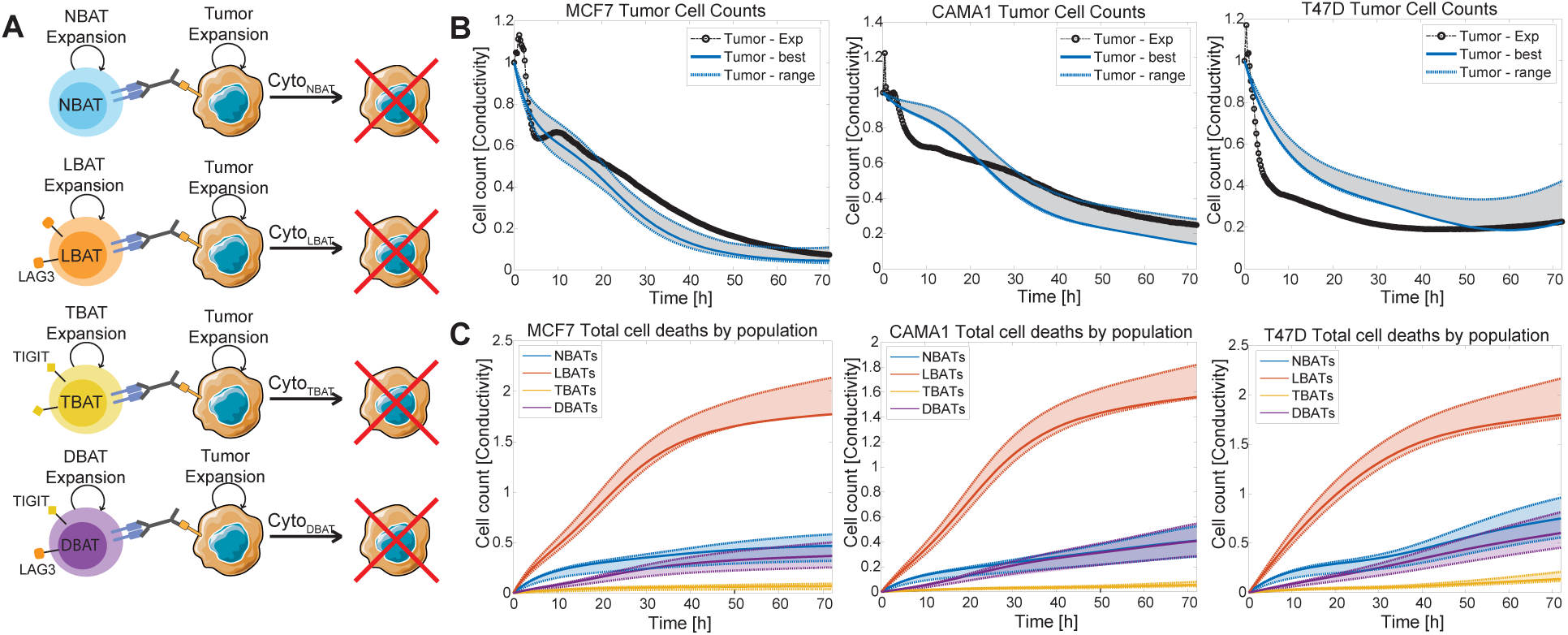
A: Diagram representing processes represented in the model. Tumor expansion parameters were determined by fitting a sigmoidal curve to tumor growth data from control conditions in which BATs were absent. BAT subgroup expansion was fit to individual splines to capture the count data determined from flow cytometry. CaliPro was used to fit cytotoxicity rates for each BAT subgroup to match the experimentally measured BAT and tumor cell counts. Data from the 3 Cancer experiment was used in parameter determination and is shown in B and C. B: MCF-7 (left), CAMA1 (center), and T47D (right) cell counts as measured by RTCA (black) and as predicted by the model (blue). Optimal model prediction in sold blue line, while range of predictions depicted by dotted blue lines. C: MCF-7 (left), CAMA1 (center), and T47D (right) cell reduction by BAT phenotype as predicted by the model. LBAT (orange), NBAT (blue), TBAT (yellow), and DBAT (purple) cumulative reduction is shown for optimal conditions (solid line) and for total range of predictions (dotted lines).

Our ODE model was constructed and distinct cytotoxicity rates were estimated for each BATs phenotype, accurately replicating tumor cell count dynamics over a 72-h BAT-tumor coculture. Tumor growth rates were determined by fitting experimental tumor growth data to sigmoidal curves, using maximum cell counts observed in tumor-only control conditions. Experimentally collected BATs counts were modeled using spline functions to represent dynami changes in BATs numbers over time. CaliPro, an iterative parameter optimization package, was employed to estimate the cytotoxicity parameter space. Value ranges for cytotoxicity rates for each BAT phenotype were identified to best match experimental BATs-tumor dynamics (**Fig 4B**). Finally, estimated cytotoxicity rates and the total number of tumor cells killed by each BATs subpopulation were compared as key metrics to evaluate the cytotoxic potential of each cell phenotype.

When comparing a single donor across 3 HER2−/HR+ breast cancer cell lines, LBATs emerge as the clear driver of cytotoxicity (**Table 2** and **Fig 4C**). LBATs demonstrate a larger capacity for tumor clearance both through a higher cytotoxicity rate (**Table 2**) and greater number of tumor cells removed compared to all other species (**Fig 4C**). This model prediction matches the functional potential observed in our PLS-R model, where LAG3+ BATs correlated with greater tumor reduction (**Fig 3B**), and our CITE-seq data, where LAG3+ CD8_Eff cells expressed significantly higher granzyme and perforin concentrations than other observed groups (**Fig 1D**). Both factors make the generation of these cells highly desirable therapeutically. Additionally, LAG3+ BATs were observed to proliferate strongly in all systems, indicating they can sustain an effective anti-tumor response when generated. DBATs, which expressed both LAG3 and TIGIT, were observed to also contribute to cytotoxicity, although at a lower rate than LBATs (**Fig 4C**). NBATs were 5.75-fold less cytotoxic than LBATs, although their presence as the most populous BAT subtype meant they contributed significantly to tumor removal. TBATs demonstrated significantly lower cytotoxicity compared to either LAG3+ subgroups LBATs or DBATs and remained at low enough numbers to contribute minimally to tumor reduction.

**Table 2:**
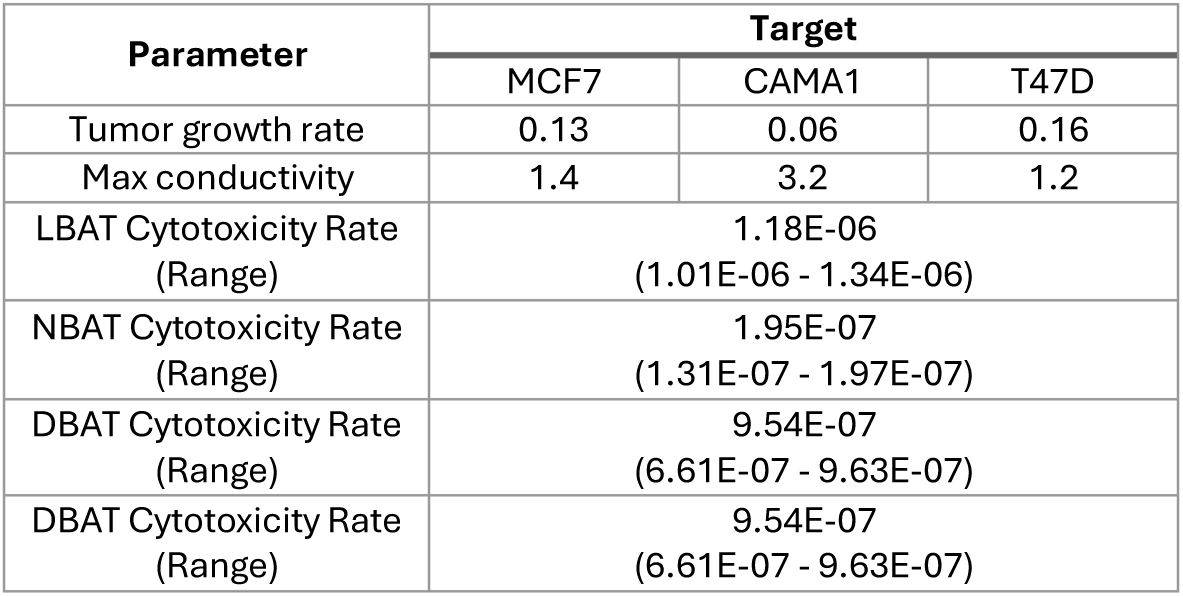
Optimal parameters and ranges determined by CaliPromeeting pass criteria for Donor 1 BATs challenged against 3 tumor cell lines.

| Parameter | Target |  |  |
| --- | --- | --- | --- |
|  | MCF7 | CAMA1 | T47D |
| Tumor growth rate | 0.13 | 0.06 | 0.16 |
| Max conductivity | 1.4 | 3.2 | 1.2 |
| LBAT Cytotoxicity Rate (Range) | 1.18E-06<br>(1.01E-06 - 1.34E-06) |  |  |
| NBAT Cytotoxicity Rate (Range) | 1.95E-07<br>(1.31E-07 - 1.97E-07) |  |  |
| DBAT Cytotoxicity Rate (Range) | 9.54E-07<br>(6.61E-07 - 9.63E-07) |  |  |
| DBAT Cytotoxicity Rate (Range) | 9.54E-07<br>(6.61E-07 - 9.63E-07) |  |  |

Comparisons across donors revealed consistent patterns of cytotoxicity. While the behavior of BATs derived from different donors could not be described using identical parameter sets, individual models predicted LBATs as the most cytotoxic subset, followed by DBATs, with NBATs and TBATs alternating as the least and second least cytotoxic subgroup (**Fig 5B-C**, **Table 3**).

**Figure 5:**
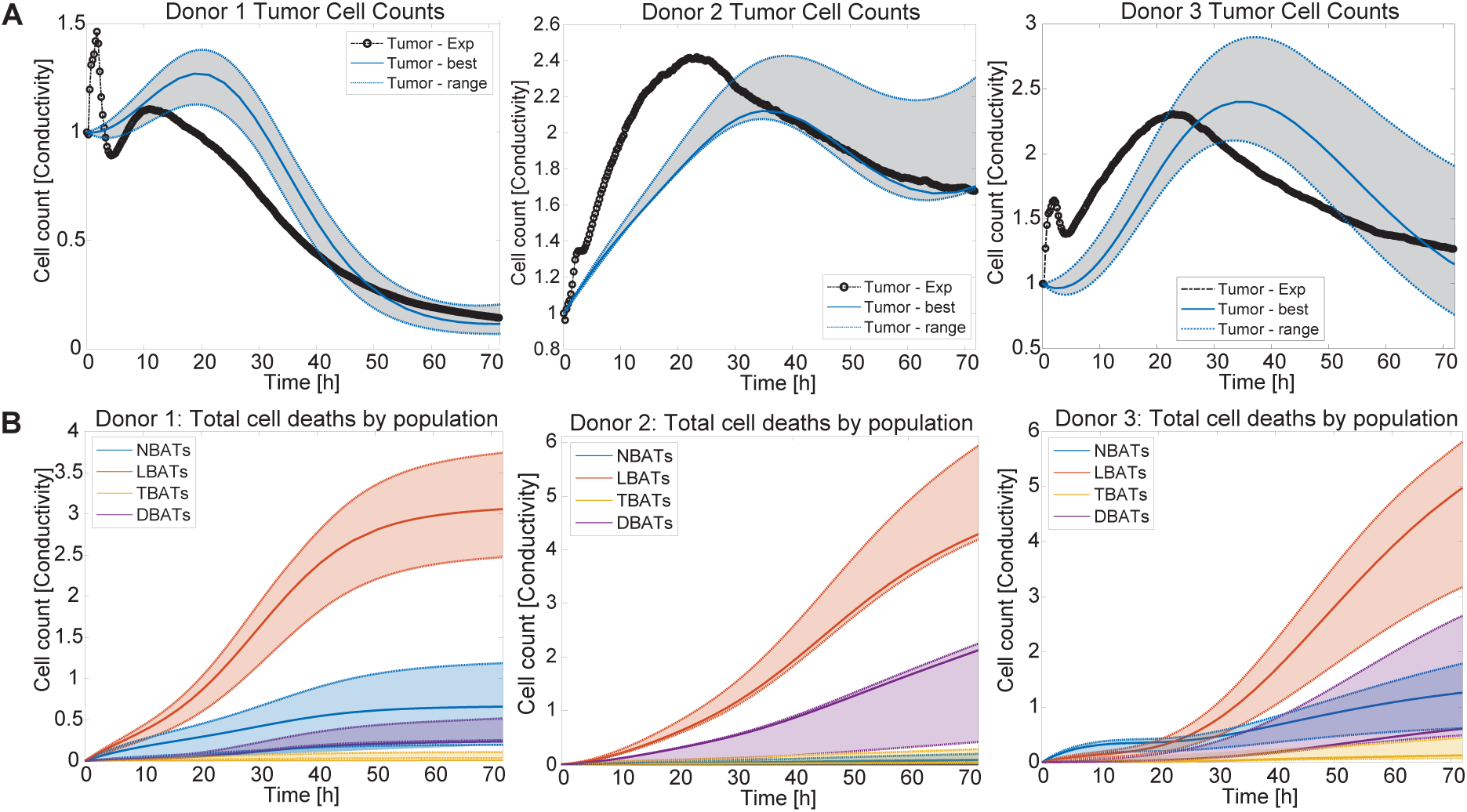
A: MCF-7 cell counts as measured by RTCA (black) and as predicted by the model (blue) when challenged by Donor 1 (left), Donor 2 (middle), and Donor 3 (right) BATs. Optimal model prediction in sold blue line, while range of predictions depicted by dotted blue lines. B: MCF-7 cell reduction by BAT phenotype as predicted by the model when challenged by Donor 1 (left), Donor 2 (middle), and Donor 3 (right) BATs. LBAT (orange), NBAT (blue), TBAT (yellow), and DBAT (purple) cumulative reduction is shown for optimal conditions (solid line) and for total range of predictions (dotted lines).

**Table 3:** Optimal parameters ranges determined by CaliPro meeting pass criteria for BATs from three donors challenged against MCF7 tumor cells.

| Parameter | Donor |  |  |
| --- | --- | --- | --- |
|  | Donor 1 | Donor 2 | Donor 3 |
| Tumor growth rate | 0.0748 |  |  |
| Max conductivity | 7.6 |  |  |
| LBAT Cytotoxicity Rate (Range) | 1.18E-06<br>(1.01E-06 - 1.34E-06) | 8.59E-06<br>(8.38E-06 - 1.16E-05) | 5.28E-06<br>(3.89E-06 - 5.95E-06) |
| NBAT Cytotoxicity Rate (Range) | 1.95E-07<br>(1.31E-07 - 1.97E-07) | 4.48E-08<br>(2.17E-08 - 4.91E-08) | 4.49E-07<br>(5.61E-08 - 7.48E-07) |
| DBAT Cytotoxicity Rate (Range) | 9.54E-07<br>(6.61E-07 - 9.63E-07) | 3.71E-06<br>(7.90E-07 - 4.35E-06) | 1.33E-06<br>(1.27E-06 - 5.52E-06) |
| TBAT Cytotoxicity Rate (Range) | 3.98E-07<br>(3.23E-07 - 4.25E-07) | 8.59E-08<br>(5.42E-08 - 5.74E-07) | 3.36E-07<br>(3.34E-07 - 7.31E-07) |

Notably, Donor 2, which demonstrated the lowest tumor cell clearance, relied more heavily on its DBATs for tumor reduction. This likely resulted from a higher relative abundance of DBATs generated during the coculture experiment compared to Donors 1 and 3.

Across all donors, LAG3 emerged as a robust indicator of effector cytotoxicity, suggesting that the LAG3+ CD8 BATs subset plays a primary role in tumor elimination during HR+ breast cancer-HER2 targeted BATs immunotherapy. While LAG3 has known inhibitory functions, it is known to be upregulated on activated cytotoxic T cells, and its presence on BATs appears to indicate a functional cytotoxic cell. In contrast, TIGIT expression in BATs cells may act either as a suppressor of cytotoxic function or an indicator of further dysfunction. To test whether the activation of either receptor was responsible for the differences in cytotoxicity observed in each species, the 3-donor experiment was repeated in the presence of varying concentrations of either an anti-LAG3 Relatlimab biosimilar or an anti-TIGIT Tiragolumab biosimilar (**Fig S9A**). The addition of anti-LAG3 generated a modest increase in cytotoxicity in Donors 1 and 3, which had higher proportions of LAG3+ CD8 BATs (**Fig S9B**). While it is not likely that MCF7 cells express LAG3 ligands MHC-II or FGL-1, LAG3 is known to be autoinhibitory, and thus the addition of LAG3 may prevent this autoinhibitory action in BATs. The addition of anti-TIGIT did not have a positive effect on BAT cytotoxicity for any donor, suggesting that TIGIT is not activated in this system (**Fig S9C**). As MCF7 cells are not known to express TIGIT ligands PVR, NECTIN1, or NECTIN2, the lack of efficacy is not surprising and suggests that TIGIT presence in our system indicates less functional BATs. Future in vivo studies should test the impact of these checkpoint blockades on BATs when in the presence of LAG3 or TIGIT ligands to confirm these results.

## DISCUSSION

Changes in receptor expression as a function of differentiation likely reflect the T cell’s capacity to mount diverse and multifunctional immune responses against perceived threats. Understanding how these phenotypes emerge, and function can significantly enhance our ability to engineer T cells for predictable and effective responses against tumors or pathogens. In this study, we characterized emerging T cell subtypes in BATs challenged by HR+ HER2− breast cancers and developed a model combining T cell inhibitory receptor expression dynamics with cytotoxicity rates to identify associations between targetable pathways of T cell inhibition and T cell anti-tumor cytotoxicity.

While inhibitory receptor expression is intrinsically linked with T cell stimulation, the factors driving differential expression remain unclear(44–46). Across three donors and when challenged with 3 HR+ breast cancer cell lines, only TIGIT, LAG3, and PD-1 were expressed within the first 72 hours of challenge. This suggests that combinatorial therapies pairing BATs with CPI therapies should prioritize these receptors.

Our model indicates an association between TIGIT surface expression and reduced BATs function. Because TIGIT ligands PVR, NECTIN1, and NECTIN2 have been detected in breast cancer and correlate with poor prognostic outcomes, the loss of function in TIGIT+ cells may result of TIGIT-ligand driven inhibition(47, 48). Blocking TIGIT may reverse the suppressive regulatory effect and augment the cytotoxicity of TIGIT+ and LAG3+/TIGIT+ subset of BATs. However, TIGIT presence on T cells may indicate a more terminally differentiated and exhausted state. Coexpression of TIGIT with other inhibitory receptors, including PD-1 and LAG3, is a hallmark of T cell dysfunction in tumors(49–52). Because we observe TIGIT expression primarily in tandem with PD-1 or LAG3, these cells may be terminally exhausted and less capable of mediating cytotoxic activity even if TIGIT activation or activity is blocked.

LAG3+ cells do not appear to be inhibited in our model. LAG3 expression in the absence of other inhibitory receptors, such as TIGIT or TIM-3, is a hallmark of cytotoxic effector T cell activation(53–55). LAG3 has been observed as primarily regulating effector functions like cytotoxicity for tumor-targeting T cells, and thus LAG3 expression by BATs may indicate a highly functional effector group(56, 57). These studies also observe PD-1 as primarily responsible for inhibiting cell proliferation, and thus the lack of PD-1 on our LAG3+ population may facilitate the proliferation observed in this population. The primary LAG3 ligand, MHC-II, is expressed predominantly on antigen-presenting cells (APCs), and thus should not be present in our coculture system(58). However, as breast cancers can be enriched in APCs, including pro-tumor macrophages, LAG3+ BATs may encounter strong inhibition when applied in vivo(59, 60). Thus, preemptive blockade of LAG3 may ensure these primary cytotoxic cells maintain their function in patients, regardless of APC presence. Additionally, LAG3 has been observed to inhibit T cell cytotoxicity without stimulation(61). Thus, LAG3 blockade may improve BATs function regardless of APC presence.

The two-cell, two-dimensional model we generated focuses exclusively on BAT changes that may be triggered through interactions with tumor cells, other T cells, or the cell signals produced by tumors and T cells. In vivo, the TME will contain a far more diverse array of cells and signals, many of which may affect BAT development and expression of LAG3, TIGIT, or related receptors. Other receptors may become more significant players in BAT function and inhibition when BATs are exposed to the wider range of signals in the TME. However, the direct tumor-BAT and BAT-BAT interactions our model accounts for will play a large role in BAT development, and thus the findings here will likely still improve BAT function.

Our investigation was limited to the dynamic changes in T cell behavior, without capturing tumor cell ligand expression dynamics. We assumed that components governing the rate of interaction between BATs and target cells remain constant, except for changing cytotoxicity observed in different BATs phenotypes. Furthermore, our coculture system was focused strictly on BiAb-mediated CD8 T cell-tumor interactions and studying signals and secreted factors from other cells was beyond the scope of our work. These factors can influence tumor and T cell behaviors affecting cytotoxicity. TCR-activated CD4 and CD8 T cells are known to produce cytokines and other signaling molecules, including interferon gamma (IFNγ) and tumor necrosis factor (TNFα)(62–64), which affect both T cell function and tumor proliferation(65), and were highly expressed in our CD8_Eff group. While the effect of these cytokines may be captured by the cytotoxicity rates predicted in our model, any compounding effects of their persistent presence or increasing concentration may have not been captured by our model. Because the goal of this project was to characterize CD8 BAT development, CD4 BATs were not included in our mechanistic model. While certainly relevant to tumor clearance, CD4 cells are largely known to be cytokine producing cells, which did not fit in our BAT-Tumor interaction model. CD4 receptor expression was largely similar across donors as well, and thus we assume that their impact is similar across experiments. Limiting the model to CD8 T cells might explain the donor-to-donor variability observed in subset-specific tumor clearance rates (**Table 3**). The initial BAT population for Donor 2 had a much higher proportion of CD4 T cells compared to Donors 1 and 3 and cleared fewer target cells (**Fig S10**). Donor 2 began with and generated fewer LAG3+ CD8s, and an increased CD4 proportion demands fewer CD8 cells, so this observation may simply be the result of fewer cytotoxic BATs present. However, CD8 BATs expansion was also lower with higher CD4, and particularly PD-1+ CD4, presence, possibly indicating an inhibitory effect elicited from CD4 BATs. The inclusion of other effector cells might further improve the model fits (**Fig 5A**) but is outside the scope of this paper.

Target cell phenotypic changes also likely change BATs-target interactions but are not captured in our system. Cytokine presence and cell damage can change receptor expression on target cells in a manner that may alter BATs-target interactions with time. Inflammatory cytokines, and in particular Type I interferons, can upregulate inhibitory receptor expression on T cells(66). Similarly, IFNγ can upregulate cognate ligands for inhibitory receptors, such as PD-L1, on tumor cells(67). Cell damage and tumor mutational status can also upregulate these ligands(68–70). Expression of PVR, a TIGIT ligand, has been observed in response to DNA damage, and may be present on some or all the target cells in our system(71). Thus, the target cells in our system may evolve to inhibit BATs expressing certain inhibitory receptors.

Preferential generation of more cytotoxic BATs subspecies may improve tumor clearance against HR+ breast cancers. Costimulatory signaling is a crucial part of T cell activation and plays a role in inhibitory receptor expression intensity(72, 73). Both our CITE-seq and flow cytometry data indicate 4-1BB expression is associated with cytotoxic effector BATs. 4-1BB was observed to be strongly correlated with LAG3+/TIGIT+ CD8_Eff cells in our CITE-seq data (**Fig 1D**), and 4-1BB ligand expression was strongly correlated with LAG3 and TIGIT expression for the initial 24-48h of our coculture experiment before disappearing from our system (**Fig 2D, Fig S6A**). This may indicate that 4-1BB exists to stimulate highly cytotoxic BATs upon their initial encounter with target cells. However, it is unclear if 4-1BB stimulation is associated with TIGIT expression alone, or if it is also linked to LAG3 expression. Further exploration of the impact of 4-1BB stimulation on BATs function and inhibitory receptor expression may provide insight into what incoming signals govern BATs function. If 4-1BB activation changes BATs function, whether to increase or decrease cytotoxicity, utilization of 4-1BB targeting therapies may enhance tumor reduction. A small number of 4-1BBL+ BATs were observed in our system at 4 hours, which may be indicative of a self-stimulating T cell subtype previously observed to increase tumor kill in mouse breast cancer models(74). However, the transient nature of both 4-1BB and 4-1BBL makes their utilization as targets difficult.

Together, these findings provide a compelling support for the use of predictive statistical analysis and mathematical modeling in the rational design of combination immunotherapies. They also strongly suggest that the anti-solid tumor activity of BATs in our phase I/II metastatic breast cancer clinical trials may be in part driven by the dominance of a LAG3+ cytotoxic phenotype over TIGIT downregulatory activity(12, 13). Our recent phase II trial in hormone refractory prostate cancer (HRPC) that combines the CPI pembrolizumab with HER2 targeted BATs showed significant clinical improvement in clinical responses and improved survival(75) when the use of CPI alone has not been successful in clinical test for HRPC. Furthermore, our recent ongoing trial combining pembrolizumab with HER2 BATs in selected metastatic breast cancer patients heavily pretreated with chemotherapy led to stable disease and improved survival (unpublished). The improvement in patient response to BATs combined with a PD-1/PD-L1 targeting therapy bodes well for further work in this area: our model predicted PD-1 to have a lower impact on BATs function than LAG3 or TIGIT and thus combining BATs with a LAG3 or TIGIT blockade may improve responses to an even greater extent. Further investigations into CPI and BATs combination strategies targeting stimulatory or inhibitory receptors to enhance BATs effector functions will likely improve the potential of BATs therapy in metastatic breast cancers and other solid tumors.

## DECLARATIONS

### Availability of data and material

Computational modeling algorithms (OPLS-R) and the mechanistic model of BAT-tumor interactions (ODE model) are available on GitHub (https://github.com/Dolatshahi-Lab). All cytotoxicity and processed flow cytometry data will be provided as Supplemental tables. Raw flow cytometry data and CITE-seq data will be provided upon request.

### Authors contributions

RWB, SD, and LGL were responsible for the conceptualization of the project and methodologies. RWB and SD wrote the manuscript with input from all authors. SD and LGL provided supervision. RWB performed experiments, analyzed data, and built the computational models. RWB, AT, SOG, LGL, and SD contributed to the design and development of the investigation. SD was responsible for funding acquisition. All authors approved the final version of the manuscript.

## Acknowledgements

This work was presented at the 2024 Society for Immunotherapy of Cancer (SITC) annual meeting (Abstract #718). We thank the Dolatshahi lab members and Dr. Kristin Anderson for providing feedback.

## LIST OF ABBREVIATIONS

TME: Tumor microenvironment
BATs: Bispecific antibody armed T cell
CPI: checkpoint inhibitor
HR+: hormone receptor-positive
LAG3: Lymphocyte activation gene 3
PD-1: Programmed Cell Death Protein 1
TIGIT: T cell immunoreceptor with immunoglobulin and ITIM domain
BiAb: bispecific antibodies
OS: overall survival
MBC: metastatic breast cancer
ODE: ordinary differential equation
ATC: unarmed activated T cells
CITE-seq: cellular indexing of transcriptomes and epitopes
RTCA: real-time cell analysis
OPLS-R: orthogonalized partial least squares regression
VIP: variable importance in projection
UMAP: uniform manifold approximation and projection
TCR: T cell receptor

