## Supplemental Figures for "LAG3+ CD8+ T cell Subset Drives HR+/HER2− Breast Cancer Reduction in Bispecific Antibody Armed Activated T Cell Therapy"

### Supplementary materials

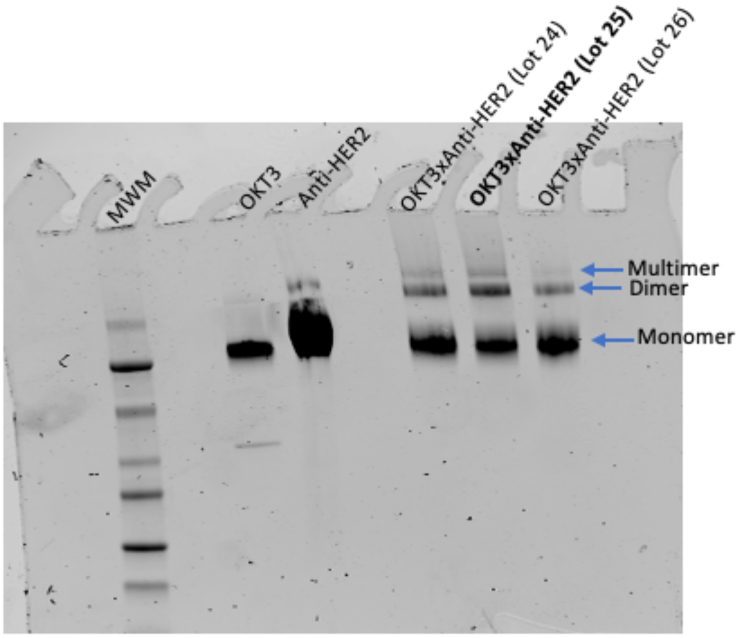


**Supplementary Figure 1:** SDS-PAGE analysis of HER2bi Lot 25 used to arm ATC in this study.

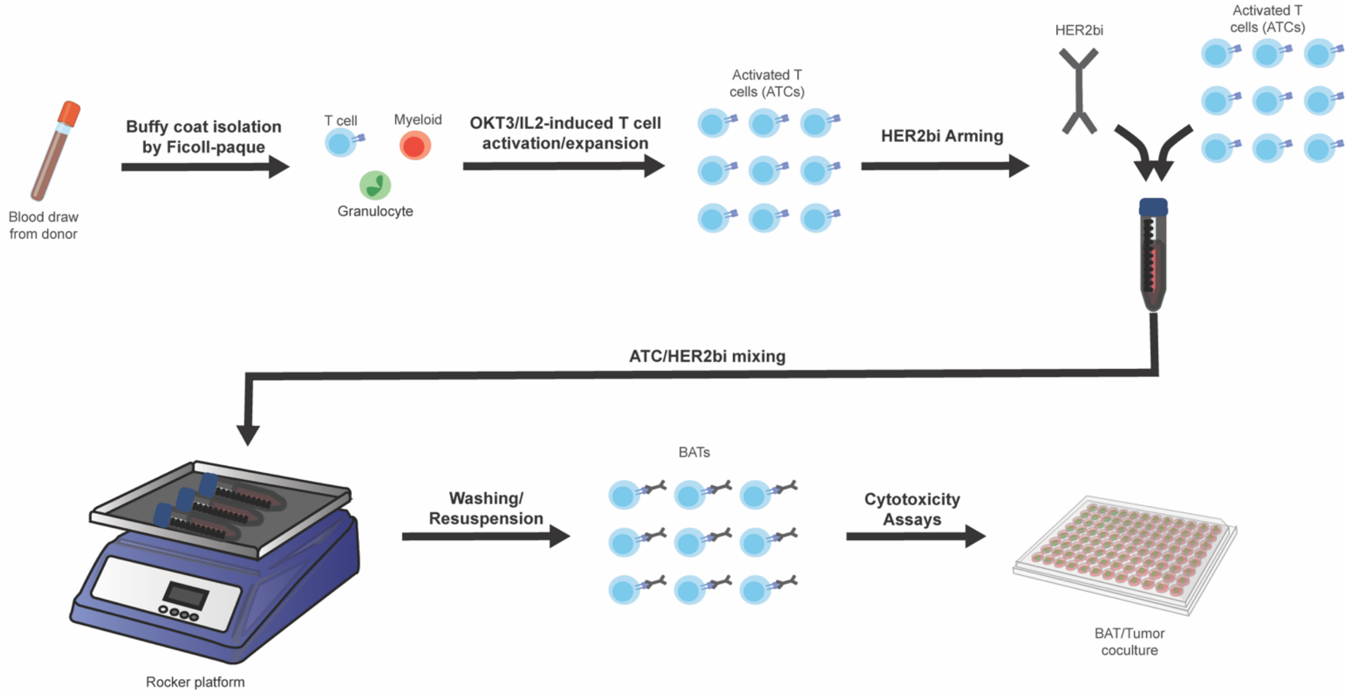


**Supplementary Figure 2:** A representative diagram of BAT production. The procedure shown was performed separately for each donor.

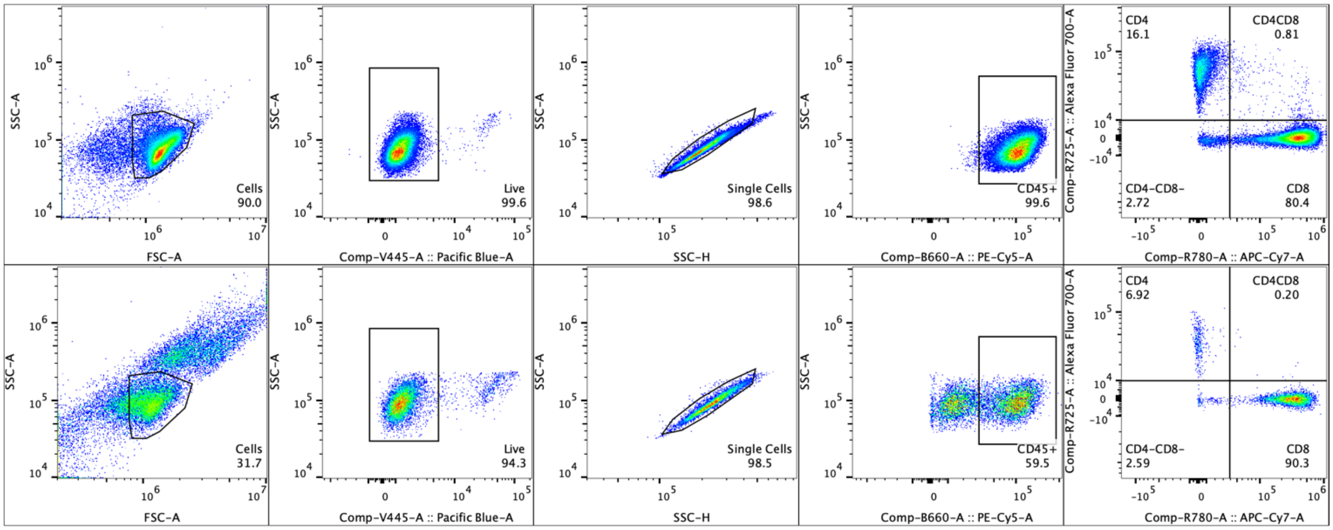


**Supplementary Figure 3:** A representative dataset demonstrating the flow cytometry gating strategy.

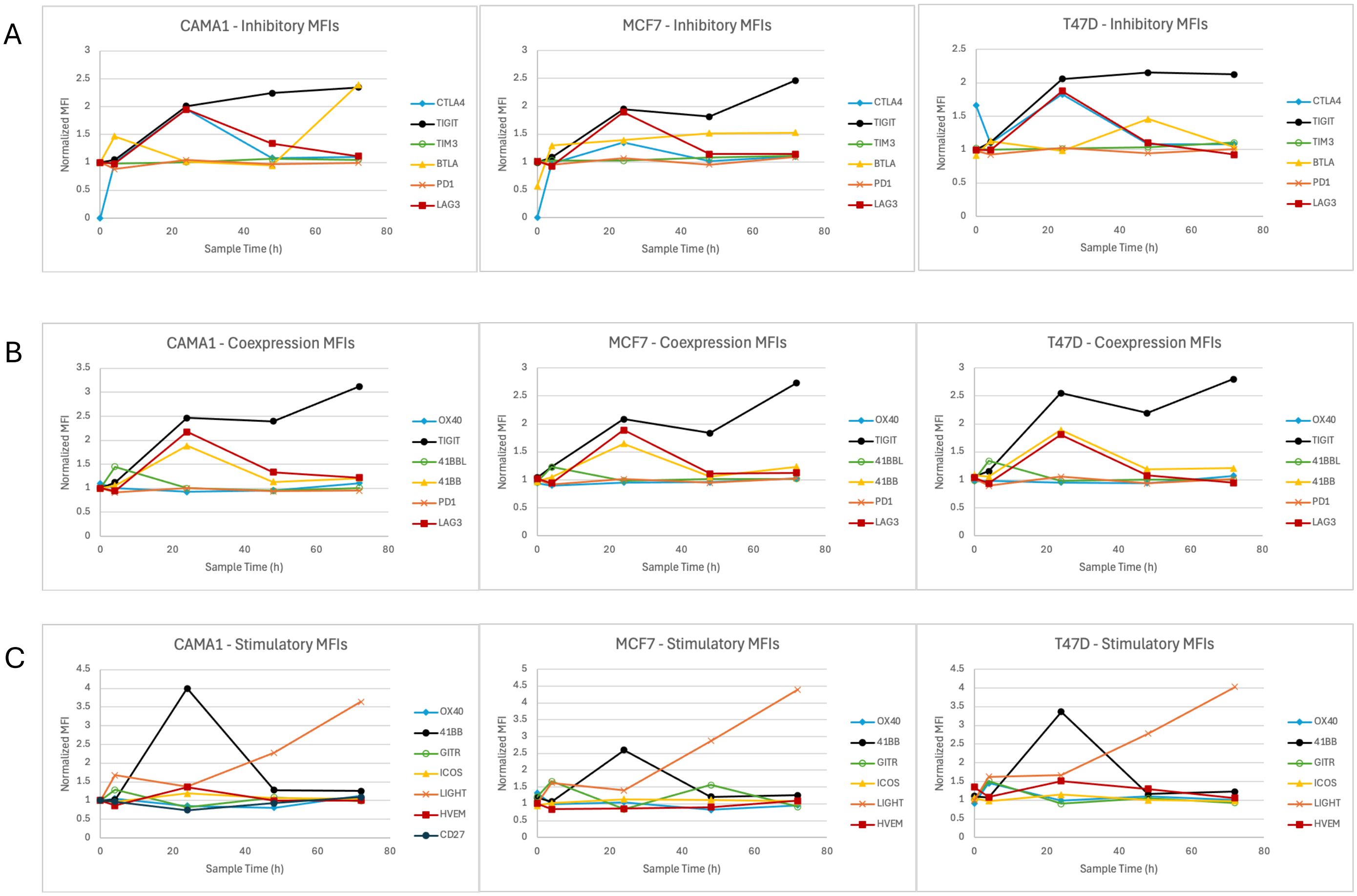


**Supplementary Figure 4:** MFI values for receptor-positive CD8 BATs from the A) Inhibitory panel, B) Coexpression panel, and C) Stimulatory panel. All MFI values are normalized to the MFI value at t=0.

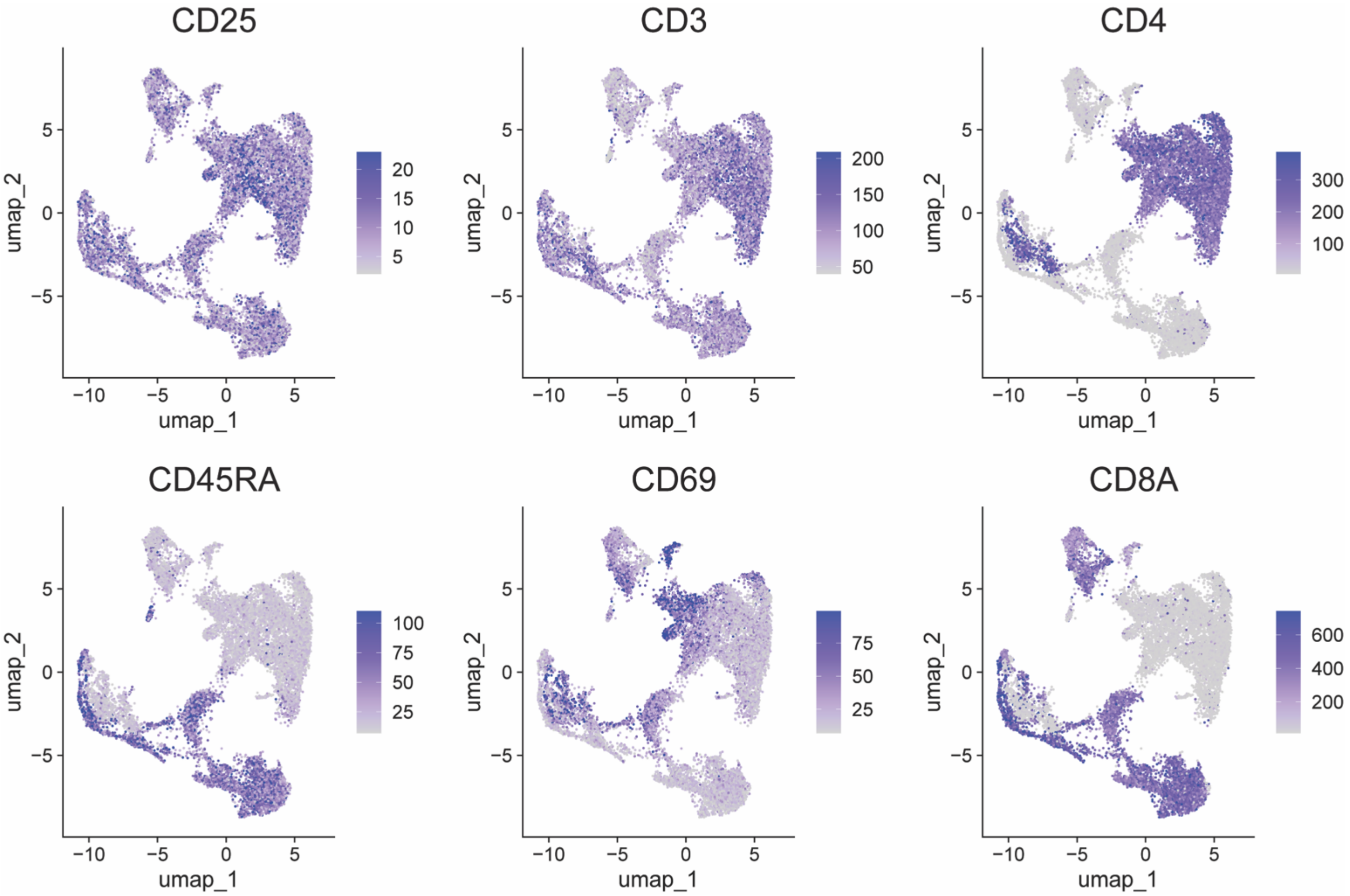


**Supplementary Figure 5:** Surface protein expression intensity of CD25, CD3, CD4, CD45RA, CD69, and CD8A measured on T cells by CITE-seq.

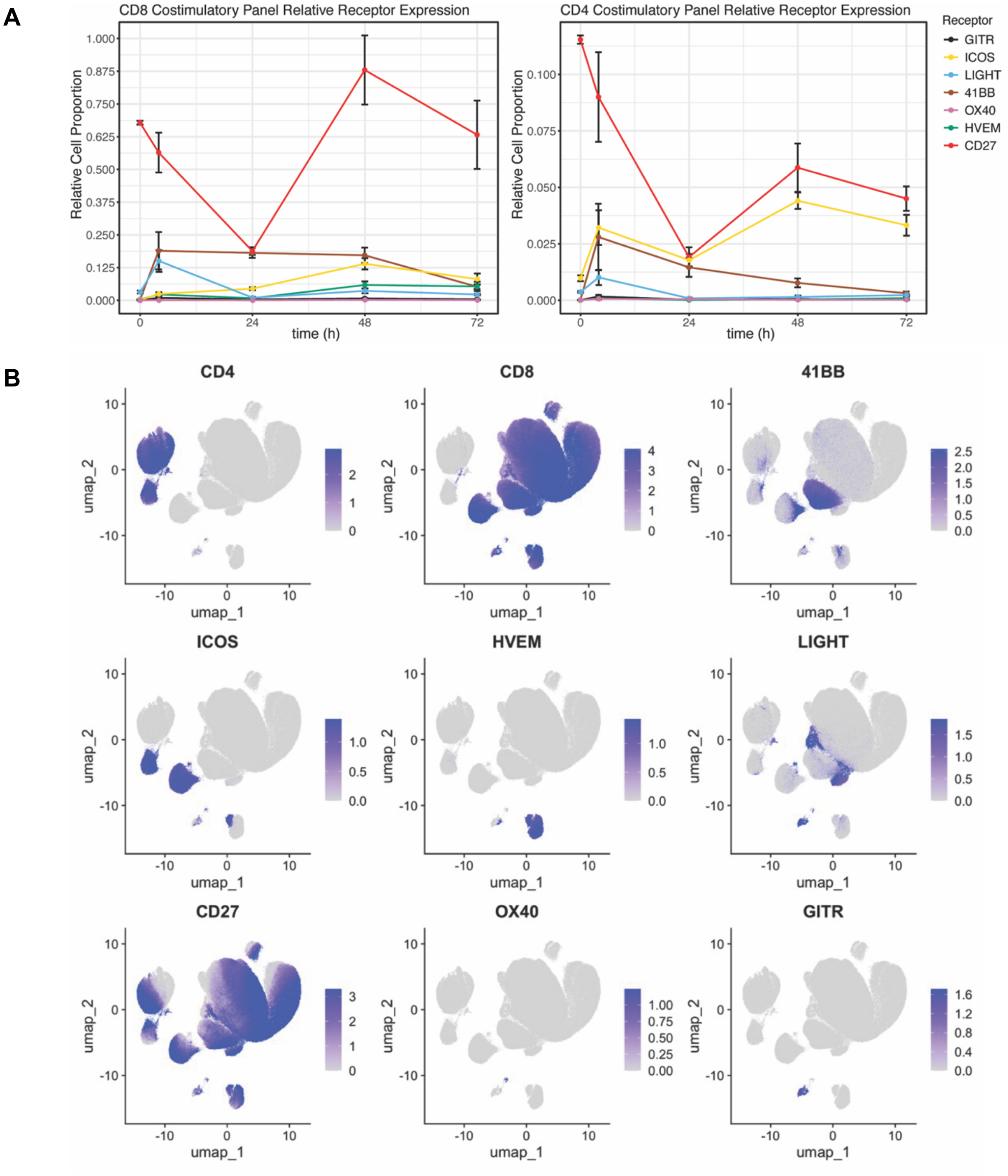


**Supplementary Figure 6:**

**A:** Supplementary panel receptor expression on CD8 (left panel) and CD4 (right panel) BATs. Data shown was collected from the Cell Line experiment and is presented as the mean value across all cell lines. Error bars indicate standard error within each data group.

**B:** UMAP of BAT cells from the Cell Line experiment clustered by receptor expression as measured by flow cytometry in the Supplementary panel.

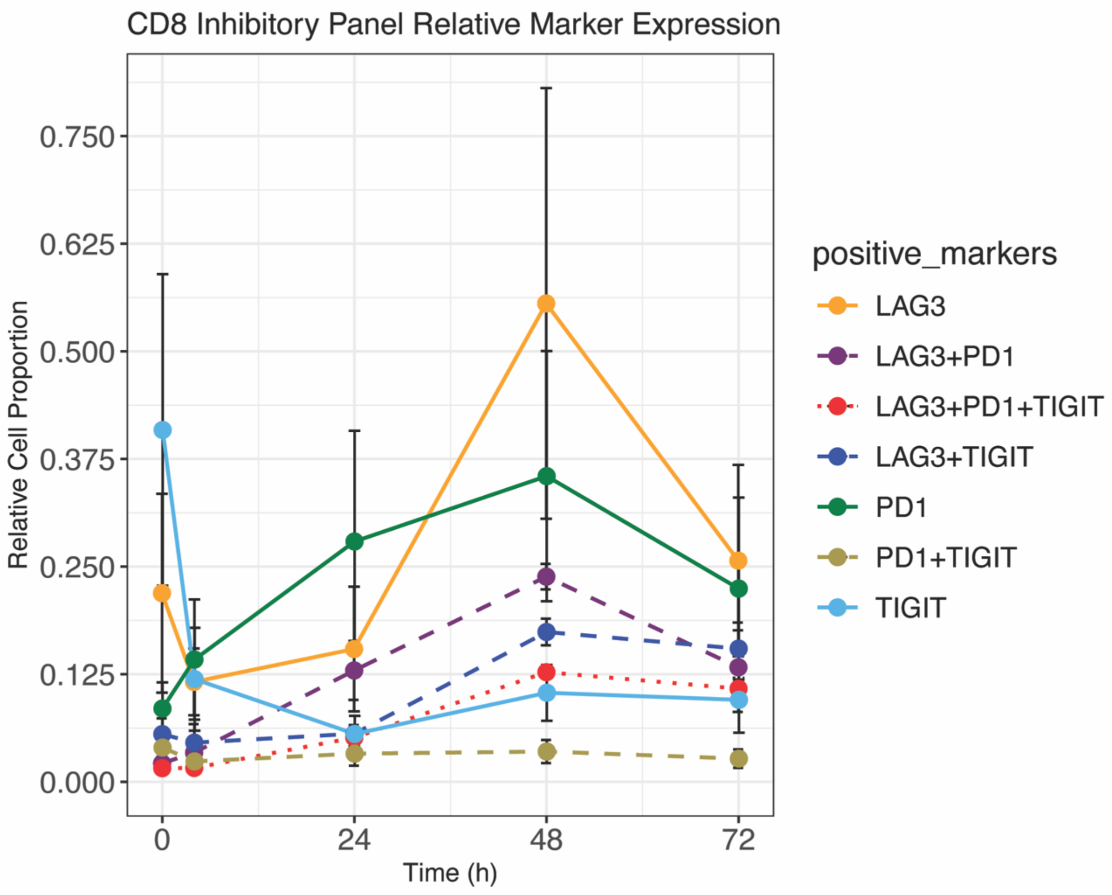


**Supplementary Figure 7:**

Relative cell proportion of CD8 cells expressing LAG3, TIGIT, PD-1, or a combination thereof. The markers that were not shown on the legend were not expressed. For example, the TIGIT+ CD8 T cell subset does not express PD-1 or LAG3. Samples collected at 0, 4, 24, 48, and 72 hours. Data represents mean value across 3 Donor experiment as collected in the coexpression panel. Error bars represent the range within each sample timepoint.

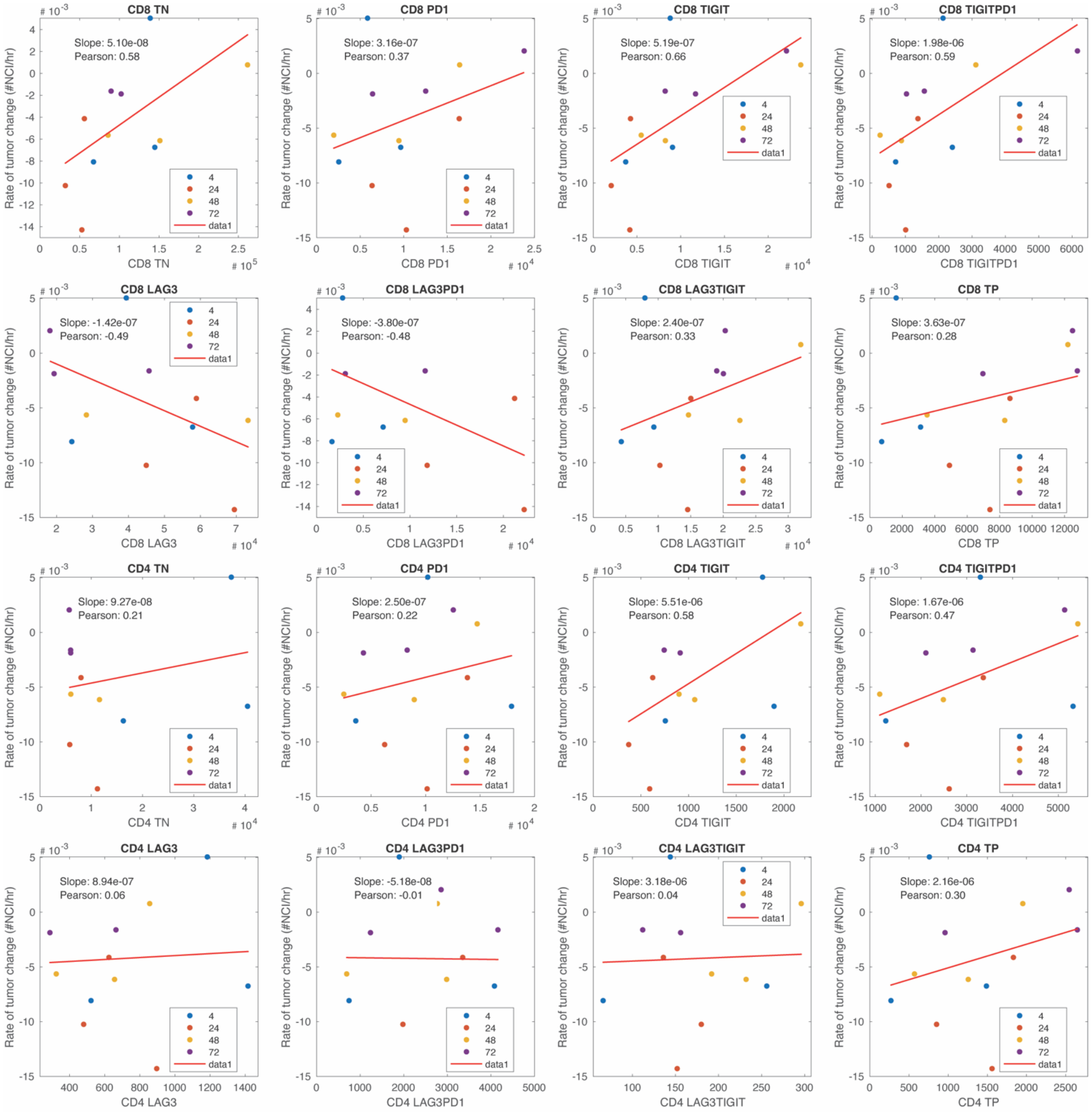


**Supplementary Figure 8:** Scatter plots comparing the viable cell count of BAT subpopulations (x-axis) to rate of tumor change (y-axis) for all subpopulations described in Figure 3B. NCI = normalized cell index.

A

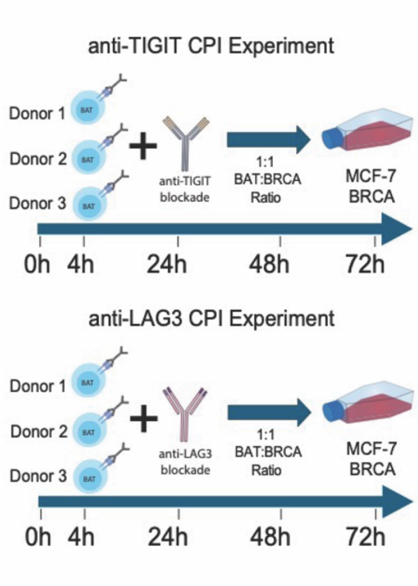


B

C

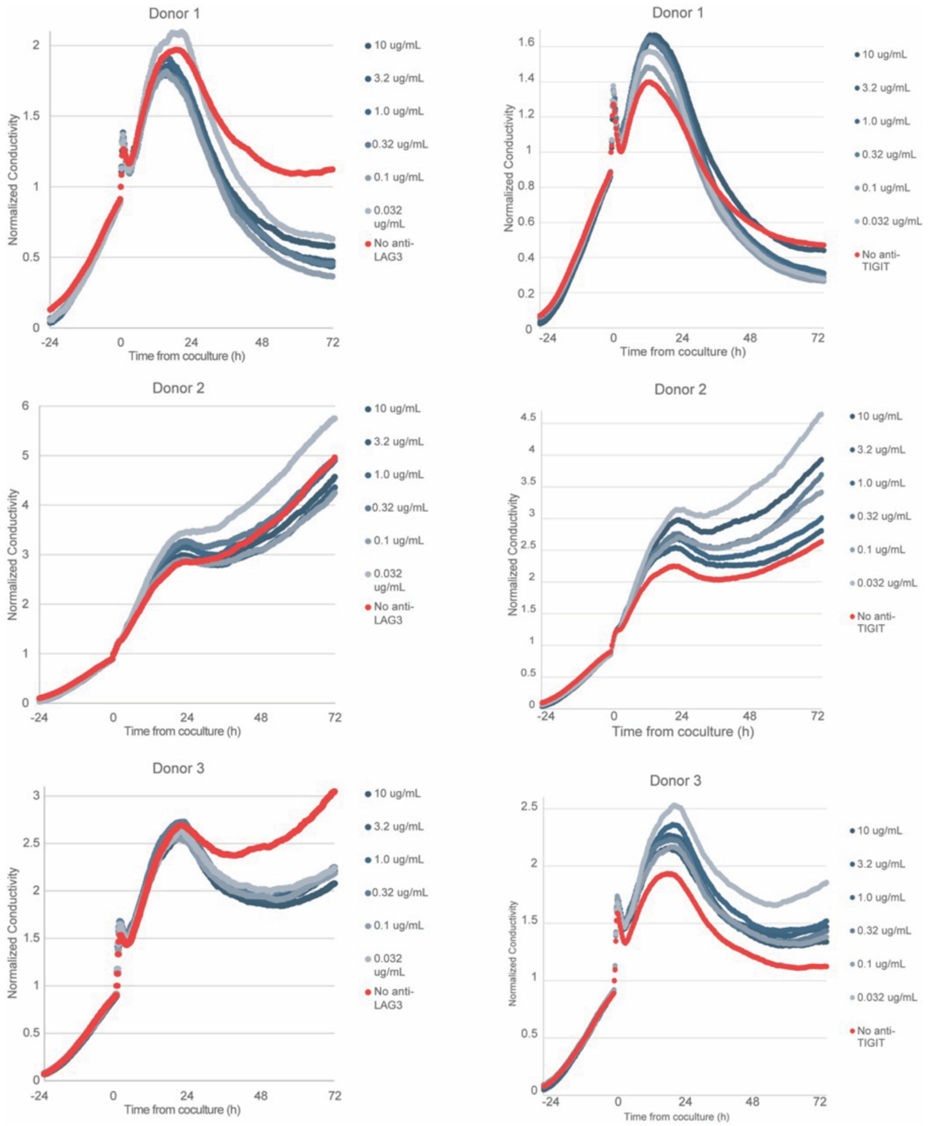


**Supplementary Figure 9:**

**A:** Pictorial representation of checkpoint inhibitor experimental setup.

**B:** Tumor cell counts for BAT-tumor coculture experiments conducted with anti-LAG3 blockade.

**C:** Tumor cell counts for BAT-tumor coculture experiments conducted with anti-TIGIT blockade.

A

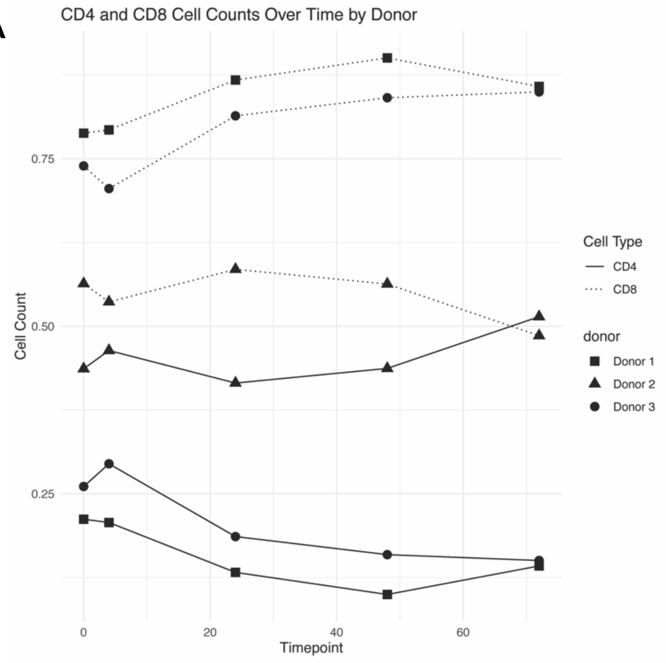


B

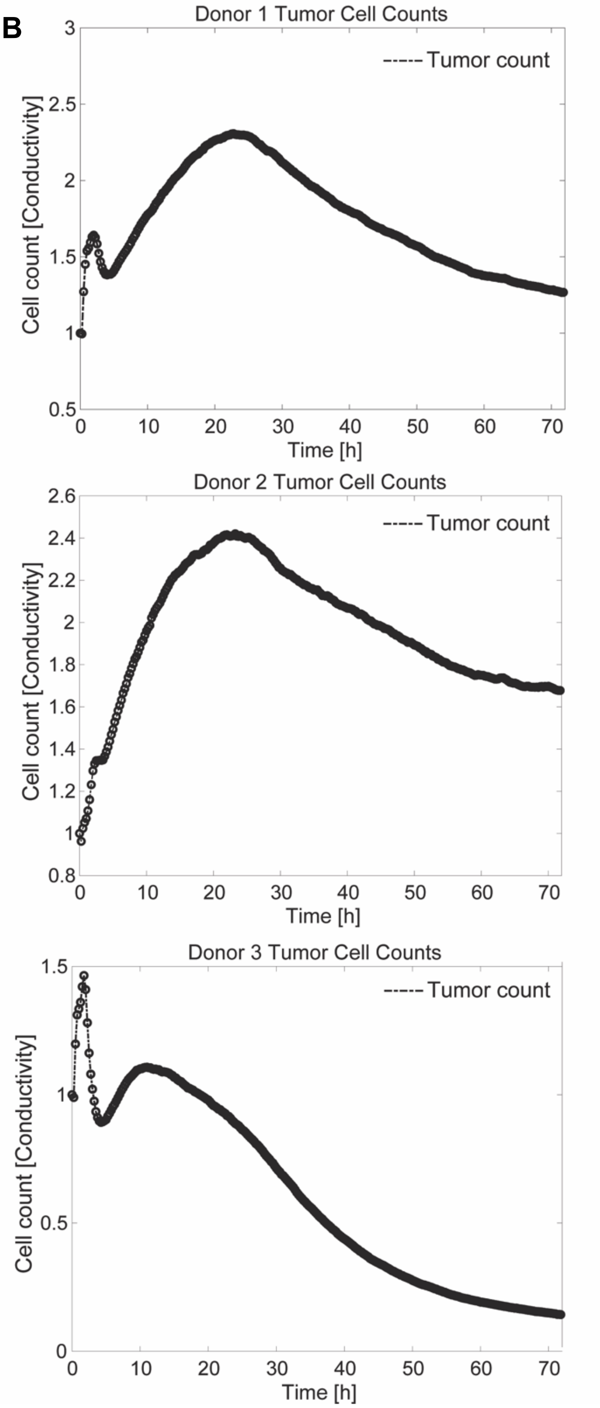


**Supplementary Figure 10:**

**A:** CD4 and CD8 proportions for each donor at time of sampling.

**B:** Tumor cell count separated by donor from the 3 Donor experiment, highlighting the donor-to-donor variability in tumor clearance.
